# Beyond commensalism: Genomic insights into micrococcin P1-producing *Staphylococcus chromogenes*

**DOI:** 10.1101/2025.08.01.668193

**Authors:** Samane Rahmdel, Tolga Türkoglu, Nastaran Nikjoo, Elham Babaali, Delara Moradi Mirhesari, Mulugeta Nega, Holger Brüggemann, Li Huang, Mathias Witte Paz, Kay Nieselt, Friedrich Götz

**Affiliations:** Microbial Genetics, Interfaculty Institute of Microbiology and Infection Medicine Tübingen (IMIT), University of Tübingen, Tübingen, Germany; Department of Food Hygiene and Quality Control, School of Nutrition and Food Sciences, Shiraz University of Medical Sciences, Shiraz, Iran; Department of Biomedicine, Aarhus University, Aarhus C, Denmark; Institute for Bioinformatics and Medical Informatics, University of Tübingen, Tübingen, Germany

**Author notes:** Corresponding authors (SR); (FG).

**Keywords:** Bacteriocin, Comparative genomics, Micrococcin P1, *Staphylococcus chromogenes*

## Abstract

*Staphylococcus chromogenes* (*S. chromogenes*) is a predominant non-aureus staphylococcal species colonizing the teat skin and mammary gland of dairy ruminants. Although often linked to mild or subclinical mastitis, specific strains may also play protective roles against major udder pathogens. In this study, we characterized two *S. chromogenes* isolates (4S77 and 4S90) that displayed antimicrobial activity against Gram-positive bacteria. Complete genome sequencing revealed a conserved, plasmid-encoded biosynthetic gene cluster for the thiopeptide bacteriocin micrococcin P1 (MP1). All genes necessary for MP1 biosynthesis, modification, export, and immunity were identified, and compound production was confirmed by HPLC and LC-MS. Comparative analysis with publicly available *S. chromogenes* genomes revealed that the MP1 cluster appears unique to these isolates. Both strains showed full phenotypic susceptibility to tested antibiotics, despite 4S90 carrying the *lnuA* gene, which did not confer detectable resistance under standard conditions. Classical staphylococcal toxin genes were also absent. Virulence gene profiling revealed a conserved repertoire of colonization- and persistence-associated genes, including factors involved in adhesion, capsule formation, and iron acquisition, but no markers of aggressive pathogenicity. Mobile genetic elements, including prophages and genomic islands, were common but did not carry antimicrobial resistance or virulence genes, suggesting a low risk of transmission of new pathogenic traits to the endogenous microbiome, including opportunistic bacteria. These findings suggest that MP1-producing *S. chromogenes* strains combine antimicrobial functionality with low virulence potential, highlighting their potential ecological role as protective commensals on the teat skin and in the broader mammary ecosystem of dairy ruminants.

## Introduction

Non-aureus staphylococci (NAS), comprising nearly 50 species, are among the most frequently isolated bacteria from the teat skin and mammary gland of dairy ruminants [1]. Widely regarded as commensals, they are also considered opportunistic intramammary pathogens, typically associated with mild or subclinical cases of mastitis. These infections often resolve spontaneously through the udder’s immune response, typically without notable impacts on milk yield or composition [2–4]. However, the contribution of NAS to overall udder health continues to be a subject of investigation, given their presence in both healthy and inflamed quarters [5]. This complexity may arise from variations among NAS species and strains in their epidemiology, virulence, pathogenic potential, ecological niches, host adaptation, and antimicrobial resistance [6]. Strains contributing to competitive exclusion and immune modulation, or producing antimicrobial compounds, are likely to be those with protective effects against major mastitis pathogens [3, 4].

*Staphylococcus chromogenes* (*S. chromogenes*) is the most prevalent NAS species predominantly isolated from milk and teat skin of dairy ruminants. Its limited presence in other environmental sources supports its classification as a host-adapted bacterium, closely associated with the udder environment [4, 6]. Strains of *S. chromogenes* differ in their capacity to induce host innate immune responses [7, 8], exhibit anti-biofilm activity against mastitis pathogens [9], and adhere to or internalize into mammary epithelial cells [10], reflecting strain-specific distinctions. In addition, various strains of *S. chromogenes* have been reported to produce antimicrobial compounds with *in vitro* activity against major mastitis pathogens, including nukacin, a cytolysin-like bacteriocin, and the purine analogue 6-thioguanine [1, 11–14]. Colonization of murine mammary glands and bovine quarters by some *S. chromogenes* strains has been shown to have a protective effect against infections by *S. aureus* and *Streptococcus uberis* [2, 4, 15]. Due to its adaptation to the ruminant mammary gland and its ability to limit infections by more pathogenic bacteria, *S. chromogenes* has been proposed as a potential probiotic for mastitis prevention and udder health enhancement [16].

In previous work, we identified antimicrobial properties in *S. chromogenes* isolates from goat and sheep milk, active against a panel of Gram-positive bacteria but not Gram-negative species [17]. In the present study, we further investigated two such isolates, 4S77 and 4S90, both recovered from goat milk, through whole genome sequencing, phenotypic profiling, and comparative genomic analysis. These strains were selected based on pulsed-field gel electrophoresis genotyping, which showed them to be the most genetically distant among eleven *S. chromogenes* isolates. We demonstrated that both strains produce micrococcin P1 (MP1), a thiopeptide bacteriocin, and described the biosynthetic gene cluster (BGC) responsible for its synthesis. Additionally, we assessed their metabolic potential, taxonomic placement, and the presence of genes associated with virulence and antimicrobial resistance, to better understand their ecological roles and potential applications in udder health management.

## Results

### General characteristics of the *de novo* assembled genomes

The circular genome maps of *S. chromogenes* strains 4S77 and 4S90 are shown in **Fig 1**. The genome of *S. chromogenes* 4S77 was assembled into a single circular chromosome of 2,318,600 bp with a GC content of 37% and one 57,477 bp circular plasmid. The genome of *S. chromogenes* 4S90 consisted of a 2,334,058 bp circular chromosome with 37% GC content and two circular plasmids of 57,607 bp and 2,638 bp. Annotation using Prokka identified 2.227 coding sequences (CDSs), 60 transfer RNA (tRNA) genes, and 19 ribosomal RNA (rRNA) genes on the chromosome of *S. chromogenes* 4S77. The chromosome of *S. chromogenes* 4S90 contained 2253 CDSs, 58 tRNA and 19 rRNA genes. The two ∼57 kb plasmids, each encoding 59 CDSs, were structurally identical, except for a 126-nucleotide difference in the length of the gene encoding lipase (GEH, glycerol ester hydrolase). The general features of ten publicly available complete genomes of *S. chromogenes* are summarised in **S1 Table**.

**Fig 1.**
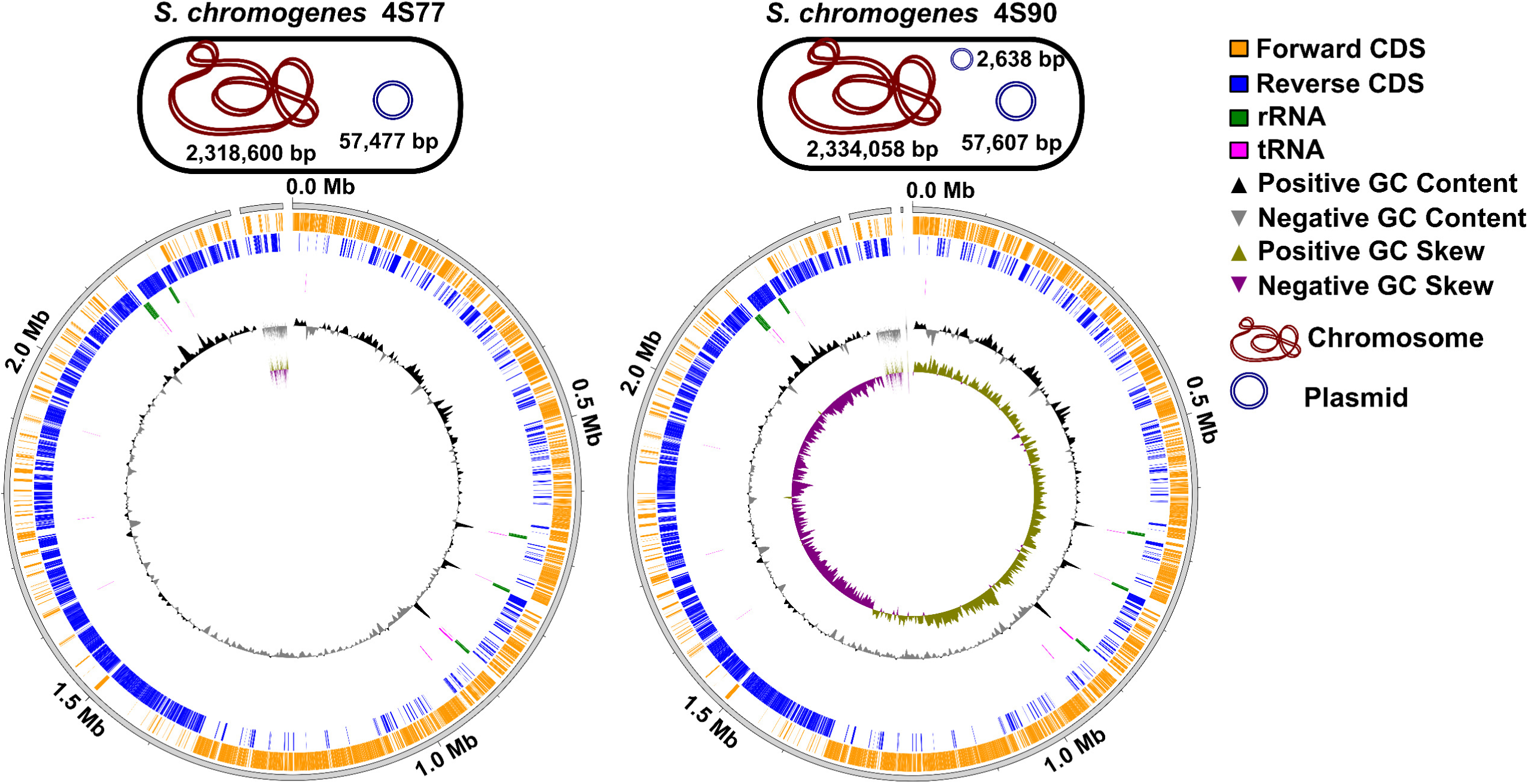
Genome map of *Staphylococcus chromogenes* strains 4S77 and 4S90.

### *S. chromogenes* phylogenetic analysis

A phylogenetic tree generated using the PATRIC (Pathosystems Resource Integration Center) tool revealed a clearly defined and well-supported clade containing all *S. chromogenes* strains, distinctly separated from other members of the genus (**Fig 2**). Within this clade, strains 4S77 and 4S90 formed a closely associated subcluster, reflecting high genomic similarity. Their placement alongside other *S. chromogenes* genomes supported their taxonomic assignment and indicated a low level of divergence from existing representatives. Notably, the *S. chromogenes* clade was positioned closest to other staphylococcal species commonly associated with veterinary sources, such as *S. delphini*, *S. felis*, *S. hyicus*, *S. muscae*, *S. schleiferi*, and *S. pseudintermedius*. The phylogenetic separation of *S. chromogenes* from other staphylococcal species was consistently robust, underscoring the discriminatory power of codon-based analysis for resolving intra- and interspecies relationships.

**Fig 2.**
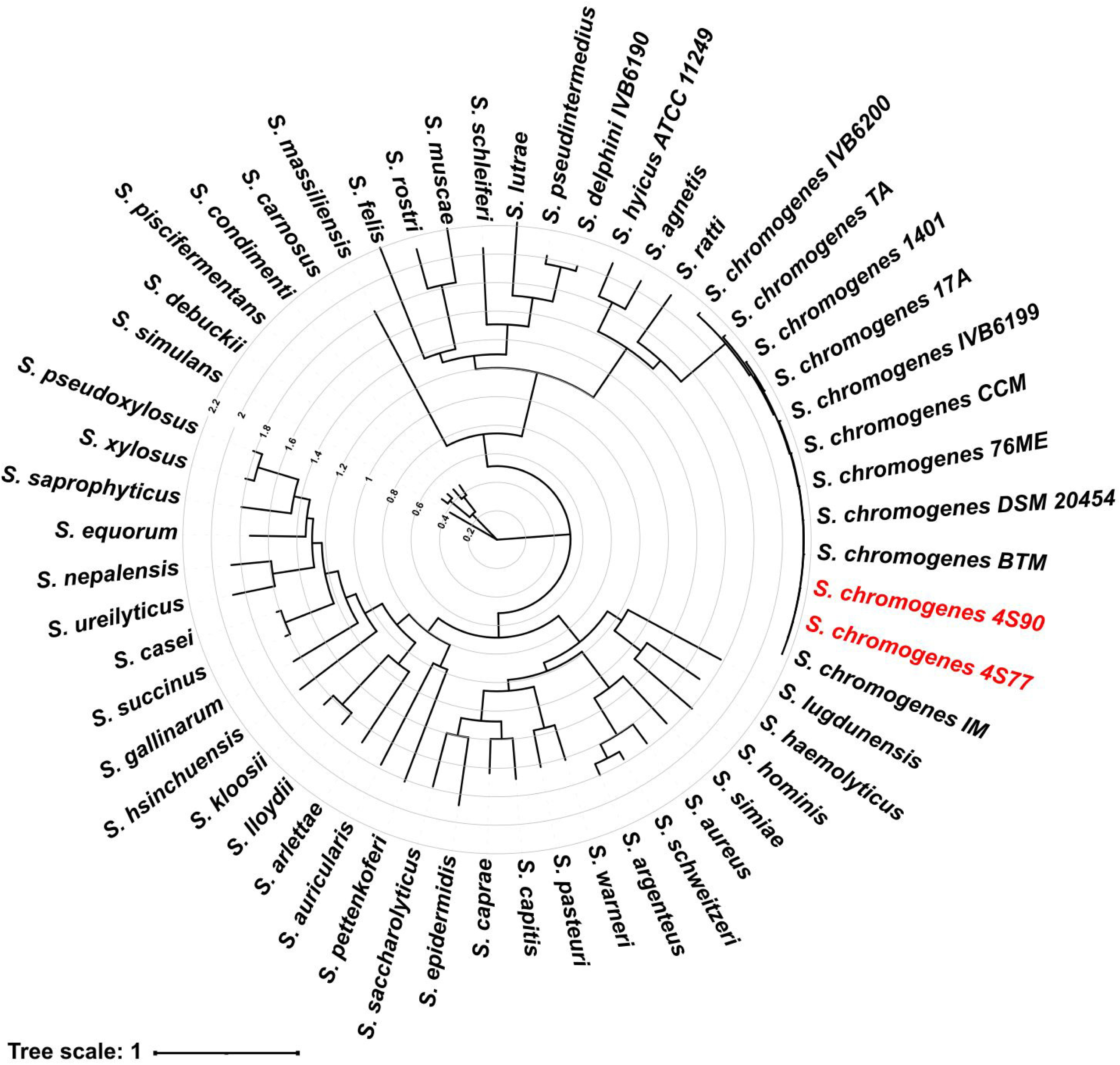
Codon-based phylogenetic tree of *Staphylococcus* species, including newly sequenced *S. chromogenes* strains 4S77 and 4S90 (this study). The tree was generated using the Codon Tree pipeline implemented in PATRIC (Pathosystems Resource Integration Center), based on alignments of 100 single-copy core genes shared across all genomes, using both amino acid and codon-aware nucleotide sequences. Alignments were performed with MUSCLE and BioPython, and phylogeny was inferred with RAxML using a partitioned model and 100 rounds of rapid bootstrapping. The analysis included 56 complete genomes: 44 reference genomes representing diverse *Staphylococcus* species, ten complete *S. chromogenes* genomes obtained from public repositories, and the two newly sequenced isolates: *S. agnetis* (GenBank accession no. GCA_011466855.1), *S. argenteus* (GCA_000236925.1), *S. arlettae* (GCA_025560845.1), *S. aureus* subsp. *aureus* (GCA_000013425.1), *S. auricularis* (GCA_016028295.1), *S. capitis* subsp. *urealyticus* (GCA_040739365.1), *S. caprae* (GCA_003966625.1), *S. carnosus* (GCA_016028275.1), *S. casei* (GCA_030294405.1), *S. chromogenes* 17A (GCA_007814525.1), *S. chromogenes* DSM 20454 (GCA_029024625.1), *S. chromogenes* 1401 (GCA_011466875.1), *S. chromogenes* IVB6199 (GCA_025558905.1), *S. chromogenes* IVB6200 (GCA_025558865.1), *S. chromogenes* 76ME (GCA_026650245.1), *S. chromogenes* CCM (GCA_048571075.1), *S. chromogenes* TA (GCA_048571095.1), *S. chromogenes* IM (GCA_048571105.1), *S. chromogenes* BTM (GCA_048571085.1), *S. chromogenes* 4S77 (this study), *S. chromogenes* 4S90 (this study), *S. cohnii* (GCA_016028115.1), *S. condimenti* (GCA_002079965.1), *S. devriesei* (GCA_007814545.1), *S. epidermidis* (GCA_000007645.1), *S. equorum* (GCA_004763965.1), *S. felis* (GCA_014133045.1), *S. fleurettii* (GCA_000420305.1), *S. gallinarum* (GCA_001593725.1), *S. haemolyticus* (GCA_000010465.1), *S. hominis* (GCA_016027875.1), *S. jettensis* (GCA_013391105.1), *S. kloosii* (GCA_016027855.1), *S. lentus* (GCA_014202265.1), *S. lugdunensis* (GCA_000025685.1), *S. massiliensis* (GCA_002849455.1), *S. microti* (GCA_000420325.1),= *S. muscae* (GCA_016028395.1), *S. nepalensis* (GCA_001592985.1), *S. pasteuri* (GCA_016028045.1), *S. petrasii* (GCA_002507345.1), *S. pettenkoferi* (GCA_016028325.1), *S. piscifermentans* (GCA_001593965.1), *S. rostri* (GCA_014202205.1), *S. saprophyticus* subsp. *saprophyticus* (GCA_000010765.1), *S. sciuri* (GCA_016027895.1), *S. simiae* (GCA_001593785.1), *S. simulans* (GCA_016028095.1), *S. stepanovicii* (GCA_014202245.1), *S. succinus* (GCA_016027655.1), *S. ureilyticus* (GCA_016027845.1), *S. vitulinus* (GCA_014202305.1), *S. warneri* (GCA_000379645.1), *S. xylosus* (GCA_016027825.1). The tree illustrates the clear separation of *S. chromogenes* from other *Staphylococcus* species, with strains 4S77 and 4S90 clustering closely within the *S. chromogenes* group. Branch lengths represent relative genomic similarity.

To refine intra-species relationships, a core-genome phylogeny was constructed using 179 *S. chromogenes* genomes (**S2 Table**), including isolates 4S77 and 4S90. The core-genome alignment covered 65% of the type strain genome (strain 17A) and provided a high-resolution set of single-nucleotide polymorphisms (SNPs) for tree construction (**S1 Fig**). Strains from diverse mammalian hosts were broadly distributed across the tree, with no distinct clades corresponding to healthy, subclinical, or clinical cases. This phylogenetic dispersion suggests substantial genomic diversity and indicates that host type or health condition does not strongly shape the evolutionary structure of *S. chromogenes* in mammals. In contrast, most chicken-derived strains clustered within a well-supported and distinct clade, pointing to a degree of host-specific adaptation in poultry that was not observed among mammalian-derived isolates.

### Metabolic pathways

Comparison of metabolic profiles revealed that all *S. chromogenes* strains, including isolates 4S77, 4S90, and publicly available genomes, formed a well-defined cluster, reflecting strong intra-species metabolic similarity. These strains consistently contained core metabolic pathways, such as glycolysis/gluconeogenesis, the pentose phosphate pathway, the citrate cycle, and essential amino acid biosynthesis. In addition, *S. chromogenes* strains exhibited higher KEGG ortholog (KO) representation and greater pathway completeness in several amino acid metabolism pathways compared to other *Staphylococcus* species, including biosynthesis of branched-chain amino acids (BCAAs including valine, leucine, and isoleucine), aromatic amino acids (AAAs including phenylalanine, tyrosine, tryptophan), as well as histidine, and lysine. They also showed consistently higher completeness in cofactor-related pathways such as pantothenate and coenzyme A (CoA) biosynthesis, and thiamine metabolism (**Fig 3**). Clustering based on pathway completeness profiles yielded a distinct and cohesive *S. chromogenes* clade, separating it from other species (**Fig 3**), indicating consistent differences in functional pathway representation across genomes. Consistent with this pattern, Jaccard similarity analysis of binary presence/absence data placed *S. chromogenes* strains in a distinct cluster together with the GRAS (generally recognized as safe) strain *S. carnosus* TM300 [18], (Rosenstein et al., 2009), and clearly distinct from *S. aureus* and *S. epidermidis* (**S2 Fig**). Among presence/absence-based methods, KO-level Jaccard analysis offered finer resolution, more effectively distinguishing species by metabolic gene content, while pathway-level clustering was more diffuse, owing to the broad conservation of core metabolic functions. These findings were further supported by an UpSet visualization, which identified a core set of 40 pathways shared across all strains, along with smaller subsets specific to *S. chromogenes* and *S. carnosus*, but absent in *S. aureus* and *S. epidermidis* (**S3 Fig**). The raw KEGG pathway annotations for each strain are available in **S3 Table**.

**Fig 3.**
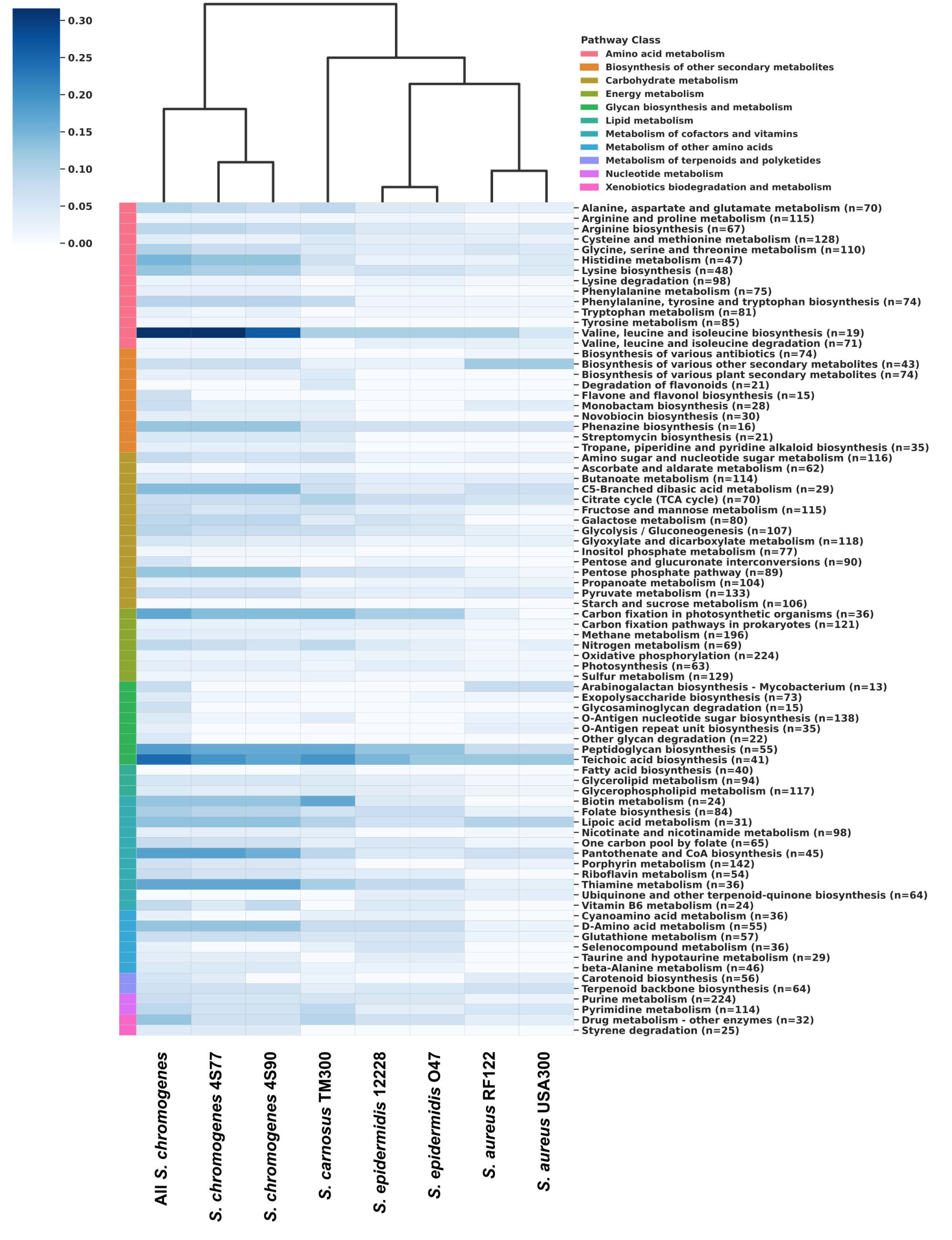
Comparative completeness of KEGG metabolic pathways of two *Staphylococcus chromogenes* isolates and reference *Staphylococcus* strains. A clustered heatmap showing the completeness of KEGG metabolic pathways across different *Staphylococcus* genomes, including two experimental *S. chromogenes* isolates (4S77 and 4S90), a composite group (“All *S. chromogenes*”) representing six reference *S. chromogenes* genomes (17A, DSM 20454, 1401, IVB6199, IVB6200, and 76ME; accessions: GCA_007814525.1, GCA_029024625.1, GCA_011466875.1, GCA_025558905.1, GCA_025558865.1, GCA_026650245.1), and reference *Staphylococcus* strains representing a range of ecological and clinical backgrounds. These include *S. aureus* USA300_FPR3757 (Human skin and soft tissue infections, GCA_000013465.1), *S. aureus* RF122 (cattle mastitis, GCA_000009005.1), *S. epidermidis* O47 (nosocomial infections, GCA_013317125.1), *S. epidermidis* ATCC 12228 (non-pathogenic, GCA_000007645.1), and *S. carnosus* TM300 (non-pathogenic, GCF_000009405.1). Completeness was defined as the proportion of annotated KO genes in each pathway relative to the total number of KO entries known for that pathway (value range: 0–1). Color intensity reflects pathway completeness (dark blue = high, light blue = low), with pathway classes indicated by a side color strip. Pathway labels indicate the name and the total number of KO entries known for that pathway. Bacterial groups were hierarchically clustered based on completeness profiles using Euclidean distance and Ward’s linkage method.

### Defense Systems

No functional CRISPR-Cas systems were detected in strains 4S77 and 4S90. Although both strains carried isolated *cas* genes and putative CRISPR arrays without adjacent *cas* loci, the high conservation of spacer sequences across *S. chromogenes* strains suggests these arrays may be false positives (**S4 and S5 Tables**). The Prokaryotic Antiviral Defence LOCator (PADLOC) analysis also revealed several shared antiviral defense mechanisms in strains 4S77 and 4S90. Both strains harbored restriction-modification (RM) Type I systems, with strain 4S90 carrying two such systems and strain 4S77 carrying one. Furthermore, both genomes included a locus containing a pseudogene (PDC S07), annotated as a “nucleoside triphosphate pyrophosphohydrolase,” as well as a Type IV RM system (**Fig 4**). Among the other analyzed *S. chromogenes* strains, the antiviral defense profiles exhibited considerable diversity, with strain-specific combinations of RM systems spanning Types I, II, and IV, CRISPR-associated elements, and other predicted phage defense systems, highlighting the complexity and variability of antiviral strategies within the species (**S6 Table**). Toxin-antitoxin (TA) loci were detected in all analyzed *S. chromogenes* strains, including 4S77, 4S90, and ten additional complete genomes retrieved from NCBI. All identified systems belonged to Type II TA modules, in which toxins are neutralized by direct binding of their cognate antitoxins. Three conserved TA systems were found across all strains: a copper-sensing transcriptional repressor/putative copper chaperone (CsoR/CsoZ), the antitoxin MazE paired with the MazF family RNA-targeting toxin, and the 6-phosphogluconolactonase/SAS053 family DNA gyrase inhibitor (**S7 Table**).

**Fig 4.**
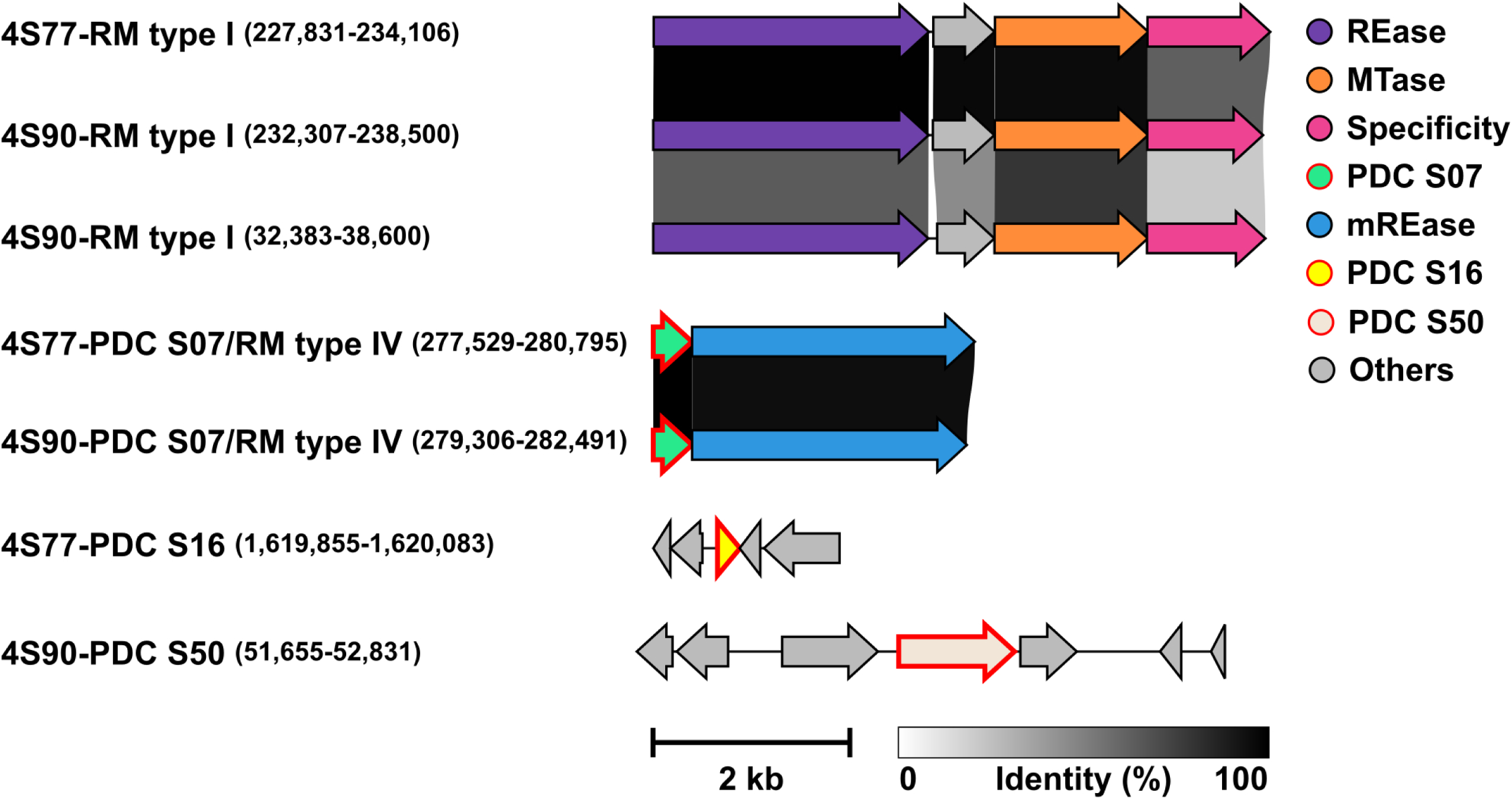
Distribution of antiviral defense systems in *Staphylococcus chromogenes* strains 4S77 and 4S90. Gene content and organisation are shown for each locus, with amino acid sequence identity between homologous genes illustrated by shaded grey connectors. Pseudogenes are indicated by red outlines. Locus coordinates are provided in parentheses after the locus name. mREase, methylated DNA restriction endonuclease; MTase, Methyltransferase; PDC, Phage defense candidate; REase, DNA restriction endonuclease; RM, restriction-modification system.

### Antimicrobial resistance (AMR) gene screening and phenotypic susceptibility

AMR gene screening across 12 *S. chromogenes* strains revealed a low overall prevalence of acquired resistance genes (**Fig 5**). The most common AMR determinant was *blaZ*, encoding beta-lactamase PC1, found in three strains (CCM, IM, IVB6200). The *lnuA* gene, associated with lincosamide resistance, was detected in two strains, including the goat-derived isolate 4S90. Additional resistance genes were found at low frequency, with *mphC* (macrolide phosphotransferase), *msrA* (macrolide-streptogramin resistance-type ABC-F protein), and *narA/B* (narasin resistance ATPase/permease) co-occurring exclusively in strain 1401, *and* str (streptomycin resistance protein) identified in a single strain (IM). Efflux pump-associated genes *sdrM* and *sepA* were detected in all strains analyzed. However, these genes were also present in the food isolate *S. carnosus* TM300 and the antibiotic-sensitive *S. aureus* strain ATCC 29213, suggesting they may represent conserved housekeeping efflux systems rather than functional resistance determinants. This interpretation was further supported by norfloxacin minimum inhibitory concentration (MIC) testing, which showed a value of 0.5 µg/mL for *S. chromogenes* strains 4S77, 4S90, DSM 20454 and *S. carnosus* TM300, indicating phenotypic susceptibility. Lincosamide susceptibility was also evaluated. Lincomycin MIC values were 0.5 µg/mL for *S. chromogenes* 4S77 and *S. carnosus* TM300, and 0.25 µg/mL for 4S90 and DSM 20454. Notably, only 4S90 carried the *lnuA* gene, raising questions about its apparent susceptibility. To further probe this, we tested clindamycin sensitivity using an MIC Test Strip; 4S90 showed an MIC of 0.125 µg/mL, confirming phenotypic susceptibility despite the presence of *lnuA*.

**Fig 5.**
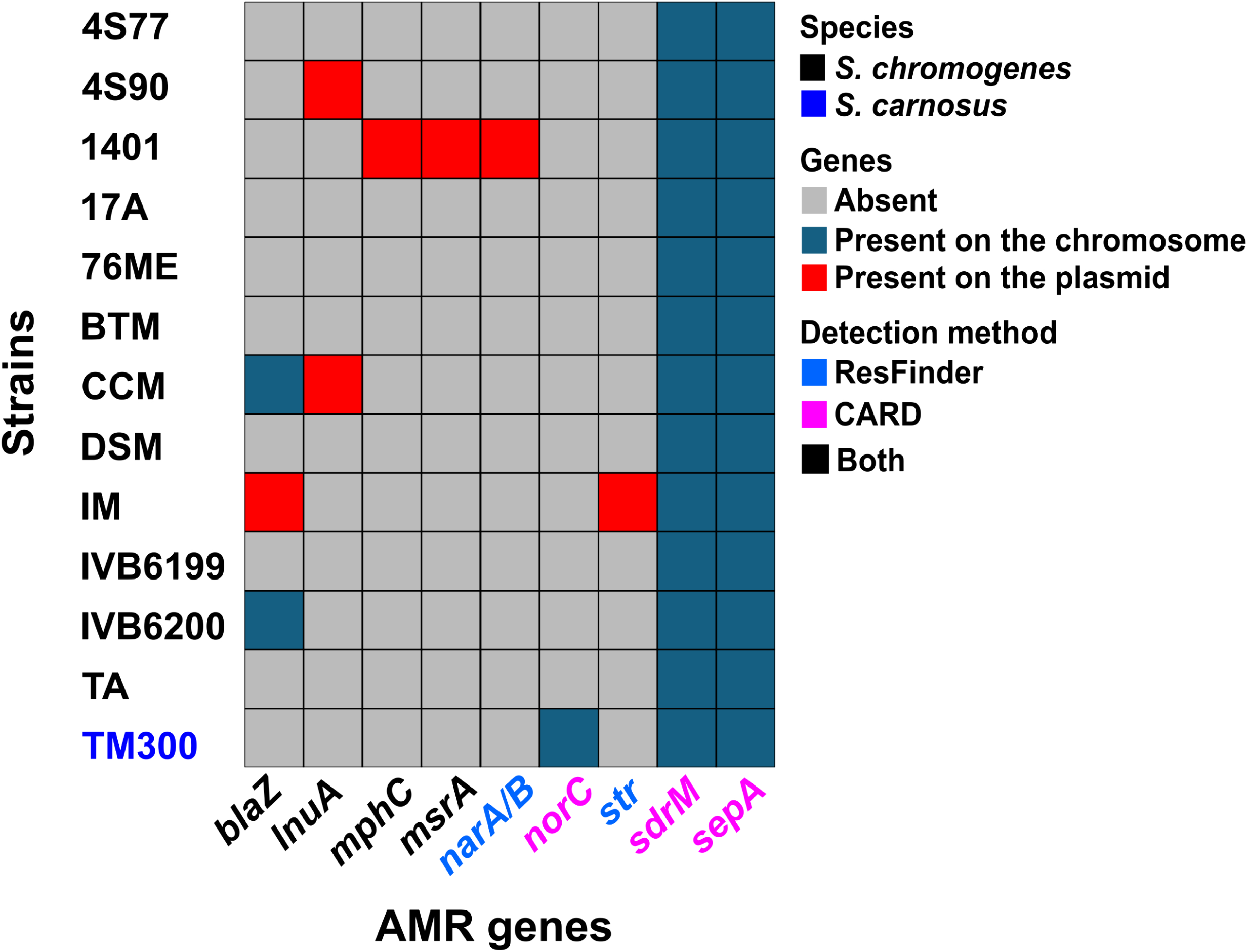
The distribution of antimicrobial resistance (AMR) genes across 12 *Staphylococcus chromogenes* strains, with *S. carnosus* TM300 included as a comparator. AMR gene colors indicate detection by ResFinder, the Comprehensive Antibiotic Resistance Database (CARD), or both tools. Strain TM300 is labeled in blue to distinguish it from *S. chromogenes* isolates. Efflux pump-associated genes *sdrM* and *sepA* were detected in all strains, suggesting they are conserved housekeeping genes rather than active resistance determinants. Consistently, all tested strains (4S77, 4S90, DSM 20454, and TM300) were susceptible to norfloxacin. Although 4S90 carried the lincosamide resistance gene *lnuA*, it remained susceptible to clindamycin, indicating limited functional impact. *blaZ*, beta-lactamase PC1; *lnuA*, lincosamide nucleotidyltransferase; *mphC*, macrolide phosphotransferase; *msrA*, msr-type ABC-F proteins; *narA/B*, narasin resistance ATPase/permease; *norC*, major facilitator superfamily (MFS) antibiotic efflux pump; *str*, streptomycin resistance protein; *sdrM*, MFS antibiotic efflux pump; *msr*, macrolide-streptogramin resistance; *sepA*, small multidrug resistance (SMR) antibiotic efflux pump. *DSM, *S. chromogenes* DSM 20454.

Alignment of the amino acid sequence of the *lnuA* gene product from 4S90 with both the CARD database reference and the homolog from *S. chromogenes* CCM revealed several amino acid substitutions (data not shown). These mutations or potentially low or absent gene expression may underlie the lack of phenotypic resistance in 4S90, despite the presence of a resistance gene. Strains 4S77 and 4S90 were previously shown to be phenotypically susceptible to a broad panel of antibiotics including gentamicin, kanamycin, penicillin G, oxacillin, vancomycin, tetracycline, erythromycin, trimethoprim, chloramphenicol, and rifampin [19], and were therefore not re-tested for these agents in the current study.

In summary, AMR genes in *S. chromogenes* were infrequent and unevenly distributed. Their functional impact appeared limited, as most strains, including both goat isolates 4S77 and 4S90, exhibited little to no phenotypic resistance under standard testing conditions.

In addition to canonical antimicrobial resistance determinants, all *S. chromogenes* strains examined possessed genetic signatures consistent with resistance to antimicrobial peptides (AMPs) (**S8 Table**). Analysis of the 12 *S. chromogenes* genomes, alongside the food isolate *S. carnosus* TM300, revealed universal presence of the *mprF* (encoding multiple peptide resistance factor) and *dltABCD* loci (responsible for teichoic acid D-alanylation), both of which mediate cell envelope modifications that reduce net surface charge and thereby diminish the binding affinity of cationic AMPs [20, 21]. All strains also encoded *vraH*, a membrane-associated protein previously implicated in AMP resistance in *S. aureus*; however, its functional role in *S. chromogenes* remains unclear, particularly given the absence of *vraDE*, with which it is known to interact to confer high-level resistance [22]. The *braSR* system, which regulates bacitracin and nisin response, was identified in 4S77, 4S90, and five other strains (BTM, IM, IVB6199, IVB6200, and TA), co-occurring with its associated ABC transporter *braDE*. However, none of the genomes contained *vraDE*, which in *S. aureus* is required for actual detoxification of these compounds. Functional studies have shown that *braSR*/*braDE* alone is insufficient to confer resistance [23], suggesting that this pathway may be non-functional in *S. chromogenes*. Crucially, none of the *S. chromogenes* genomes encoded the *graRS/vraFG* module, the only staphylococcal AMP sensor-transporter system known to induce adaptive resistance to host-derived AMPs. This system was identified solely in *S. carnosus* TM300. The absence of this pathway may limit the capacity of *S. chromogenes* to mount inducible responses to AMP exposure, leaving the species reliant on constitutive or basal defences. Genes required for poly-N-acetylglucosamine biosynthesis (*icaADBC*), which facilitates AMP repulsion in biofilm matrices [24], were uniformly absent from all genomes analysed. In contrast, the *capBCAD* operon responsible for production of poly-γ-glutamic acid (PGA), a polymer capable of sequestering both cationic and anionic peptides [24], was conserved across all *S. chromogenes* strains. The metalloprotease gene *aur* (encoding aureolysin), implicated in AMP degradation [24], was likewise present in all *S. chromogenes* genomes but absent from *S. carnosus* TM300. Neither *sspA* nor *sak*, encoding additional peptide-inactivating proteases (a serine protease and staphylokinase, respectively) [24], were detected in any of the strains.

### Virulence factors (VFs)

Virulence gene profiling of 12 *S. chromogenes* strains, including two isolates from healthy goats (4S77 and 4S90) and ten publicly available genomes, revealed a consistent presence of genes associated with adherence, biofilm formation, exoenzyme production, capsule biosynthesis, and iron acquisition (**Fig 6**). Adherence-related genes such as *atl* (autolysin) and *fnb* (fibronectin-binding protein) were detected in all strains, and also found in *S. carnosus* TM300, suggesting their likely role in adhesion and colonization. The *bap* gene (biofilm-associated surface protein) was present in 11 out of 12 strains, including goat-derived strains 4S77 and 4S90. However, all strains lacked the *ica* operon, which encodes polysaccharide intercellular adhesin [25], and the *aap* gene, which encodes the accumulation-associated protein, two key factors involved in biofilm formation. Genes encoding the exoenzymes *nuc* (thermonuclease), *aur*, and *vWbp* (von Willebrand factor-binding protein) were consistently present in all *S. chromogenes* genomes. Notably, *aur*, also discussed in the context of AMP degradation, was absent from *S. carnosus*, which carried only *nuc*. Capsular biosynthesis genes also showed a structured pattern. The *capBCAD* operon, previously noted for its role in AMP sequestration, was detected in all *S. chromogenes* strains, supporting a potential dual function in both host immune evasion and surface protection. In contrast, *Staphylococcus*-type capsular polysaccharide genes *capO* and *capP* were universally present, while other *cap* genes (*capA–capM*) varied across strains. Iron acquisition systems were likewise well conserved. All *S. chromogenes* strains encoded the staphyloferrin A biosynthetic genes (*sfaABCD*), the adjacent transporter (*htsABC*), and a homolog of *fhuC* (ferric hydroxamate uptake ATPase), which is essential for siderophore import [26]; this genomic arrangement was also conserved in *S. carnosus*. Importantly, no classical toxin genes were detected among the *S. chromogenes* strains. Genes encoding alpha-, delta-, or gamma-hemolysins (*hla*, *hld*, *hlgABC*), leukocidins/leukotoxins, exfoliative toxins, enterotoxins, and toxic shock syndrome toxin (*tsst*) were entirely absent. The only hemolysin gene consistently detected outside regulatory regions was *hlb* (beta-hemolysin), present in all *S. chromogenes* strains but absent from *S. carnosus*. In contrast, *hld* (delta-hemolysin), located within the *RNAIII* locus, was found in all *S. chromogenes* strains as well as in *S. carnosus* TM300. While TM300 also harbored *psmβ1* and *psmβ2*, only one *S. chromogenes* strain (4S90) carried *psmβ4*. Additionally, a Type VII secretion system locus was identified exclusively in strain IVB6200, highlighting further genomic divergence. A summary of virulence factors absent from *S. chromogenes* strains 4S77 and 4S90 is provided in **S9 Table.**

**Fig 6.**
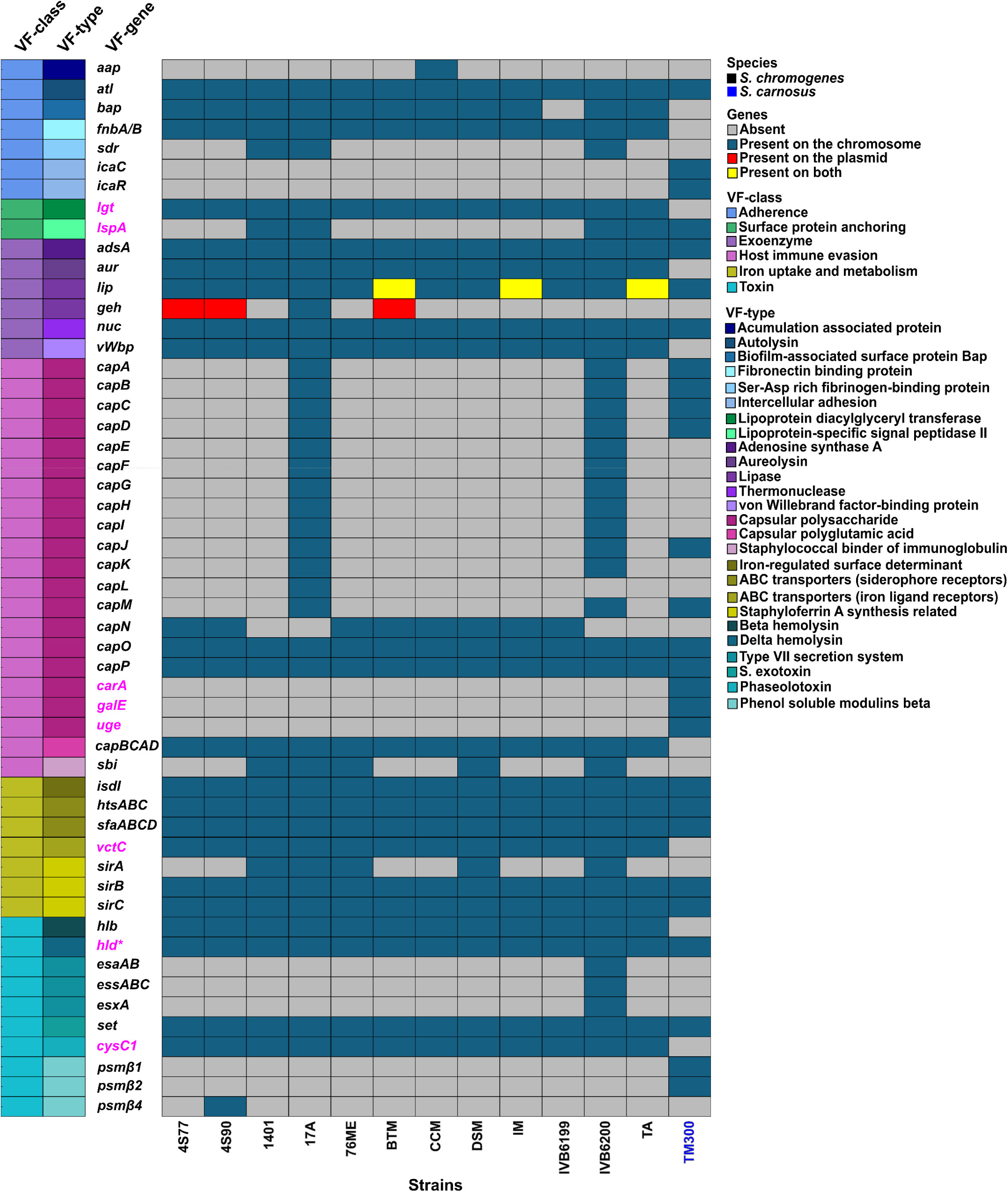
The distribution of virulence-associated genes across 12 *Staphylococcus chromogenes* strains, with *S. carnosus* TM300 included as a non-pathogenic comparator. Virulence genes were identified using BLASTx and VFanalyzer tools; genes detected exclusively by VFanalyzer are highlighted in pink. Gene presence and genomic location are color-coded as indicated: absent, chromosomal, plasmid, or present on both. Virulence factors (VFs) are grouped and color-coded by VF class and VF type on the left. Strain TM300 is labeled in blue to distinguish it from *S. chromogenes* isolates. DSM, *S. chromogenes* DSM 20454. * h*ld* was identified in all *S. chromogenes* strains through manual genome mining.

Notably, strains isolated from clinical conditions such as mastitis and intramammary infection showed no clear difference in virulence gene content compared to strains from healthy hosts. This uniformity across sources reinforces the interpretation that *S. chromogenes* harbors a conserved repertoire of colonization- and survival-associated genes, with limited evidence for classical virulence determinants. The presence of many so-called virulence genes in *S. carnosus*, a food-grade non-pathogenic species, further highlights the need for cautious interpretation of gene presence alone. Overall, the genomic evidence supports the view that *S. chromogenes*, while equipped for host interaction and persistence, lacks the toxin arsenal typical of aggressive pathogens.

This interpretation was further supported by phenotypic assays evaluating nuclease, hemolytic, lipase, and biofilm activity in *S. chromogenes* strains 4S77, 4S90, and DSM 20454 (**Fig 7**). All three strains showed functional expression of these enzymes, yet consistently at lower levels than *S. aureus* JE2. On DNase agar, *S. chromogenes* strains produced visible clearing zones, indicating nuclease activity, though these were markedly smaller than those of JE2 and absent in *S. carnosus* TM300. On sheep blood agar, the *S. chromogenes* strains displayed weak hemolysis, consistent with the presence of *hlb*, in contrast to the strong hemolysis exhibited by JE2 and the absence of hemolysis in TM300. Lipase activity on tween agar was comparable to TM300 but again fell short of the more robust activity seen in *S. aureus*. In the biofilm assay, all *S. chromogenes* strains formed dark colonies on Congo red agar, consistent with biofilm production, similar to the positive control *S. aureus* MW2. No pigmentation was observed for *S. carnosus* TM300 (**Fig 7**).

**Fig 7.**
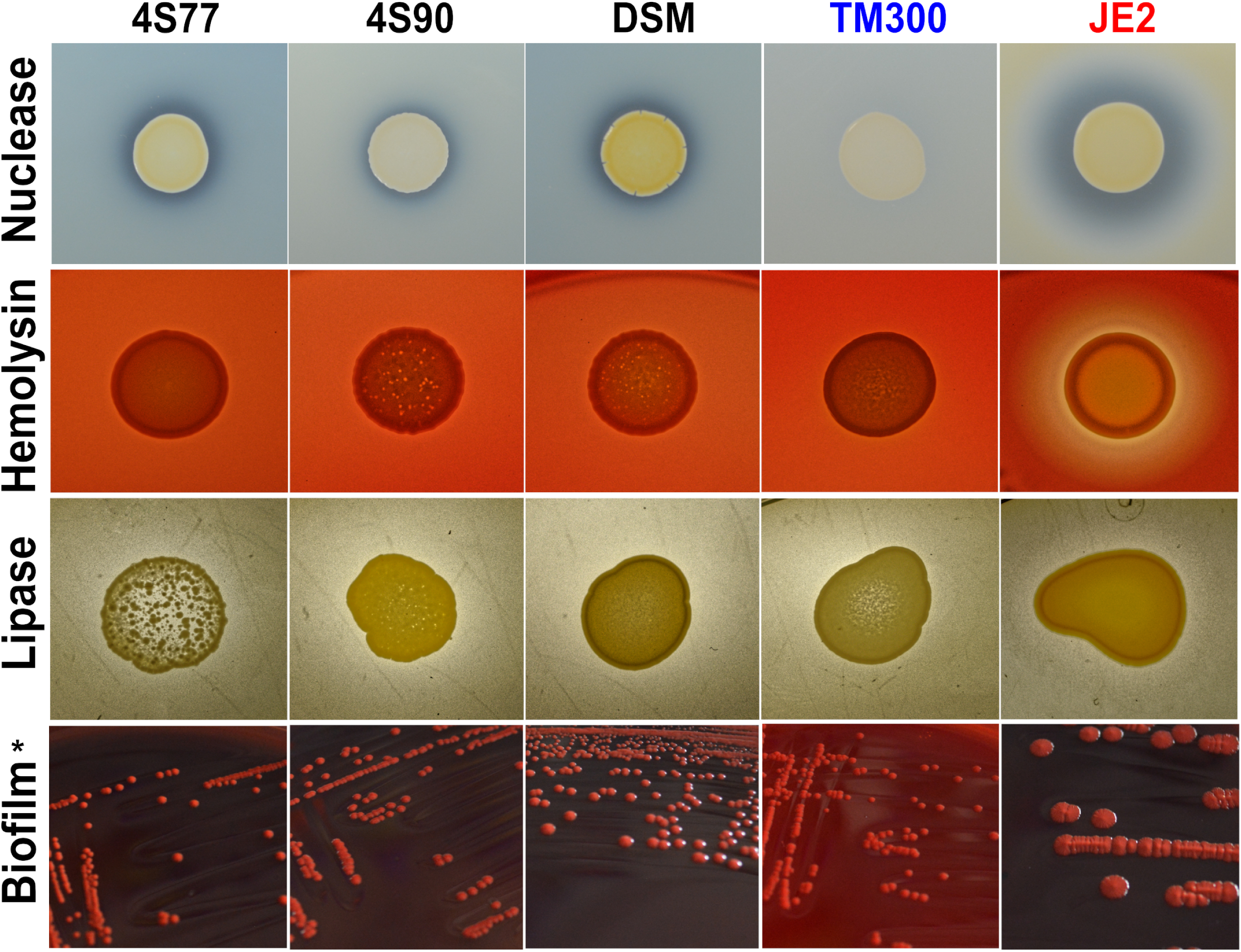
Nuclease, hemolysis, lipase activity, and biofilm formation in *Staphylococcus chromogenes* (4S77, 4S90, DSM 20454), *S. carnosus* TM300, and *S. aureus* strains. * *S. aureus* JE2 was used as the positive control in all assays except for biofilm formation, where *S. aureus* MW2 was used instead.

### Two-component regulatory systems

To explore the regulatory landscape of *S. chromogenes*, we systematically profiled two-component systems (TCSs) across 12 genomes and compared them to well-characterized systems in *S. aureus* [27]. As described above, *graRS/vraFG* and *vraDE,* key mediators of AMP resistance in *S. aureus*, were absent from all *S. chromogenes* strains, and although the *braSR*/*braDE* system was present in seven genomes, it likely lacks function due to the absence of *vraDE*. This analysis was extended to the full TCS repertoire.

Nine conserved two-component systems were detected across all 12 *S. chromogenes* genomes: *walRKHI* (cell division), *nreGYJI/nreABC* (nitrogen respiration), *hrtAB/hssRS* (heme detoxification), *RNAIII*/*agrBDCA* (quorum sensing), *vraUTSR* (cell wall stress resistance), *airSR* (redox sensing), *phoPR* (phosphate homeostasis), *srrAB* (oxidative stress), and *arlRS* (surface protein regulation and manganese homeostasis). The systems *saeRS* (toxin regulation), *lytSR/lrgAB* (autolysis and cell death), and *hptASR/uhpT* (glucose-6-phosphate import) were absent from all strains. The potassium-sensing system *kdpABC/kdpDE* (K⁺ homeostasis) was present only in strains 17A and IVB6200.

A distinct four-gene operon encoding a response regulator, ABC transporter ATPase, permease, and histidine kinase was identified in eight strains but absent from 4S77, DSM 20454, IM, and IVB6200. While the operon superficially resembled TCS–ABC transporter modules involved in peptide sensing and lantibiotic (gallidermin) resistance, its components showed no significant similarity to any of the Cpr-and Bce-type systems reported in *Clostridium difficile*, *Streptococcus mutans*, *Streptococcus agalactiae, Bacillus subtilis*, and *S. aureus* [28]. Furthermore, unlike these reference systems in which the histidine kinase and response regulator are encoded adjacently, the regulatory components in the *S. chromogenes* module were separated by the ABC transporter genes. A co-localized amidohydrolase gene was also present upstream but occurred independently in strains lacking the operon, suggesting it is not functionally linked.

Moreover, all strains harbored three unclassified TCSs with histidine kinases shared less than 30% identity to known systems, possibly representing novel or lineage-specific regulators.

### Mobile genetic elements

#### Insertion sequences (ISs) and transposons (Tns)

ISs from the IS6 family were identified on both the chromosomes and plasmids of *S. chromogenes* strains 4S77 and 4S90 (**S10 Table**). Each chromosome contained a single IS6 element with a truncated transposase, likely rendering it inactive. However, it may potentially rely on a downstream IS1182 family transposase to facilitate its movement through a hitchhiking mechanism. On the ∼57 kb plasmids of both strains, five IS6 elements were found; three had intact transposases, while the others were defective due to premature translation termination codons, potentially impairing their functionality [29]. No composite Tns were detected. However, a targeted search for Tn3-family elements identified a resolvase gene on the ∼57 kb plasmids of both strains (FDAOAELG_02344 and MNHDEDAN_02369), with no adjacent transposase detected. On the chromosome of 4S90, a DUF536 domain-containing protein (MNHDEDAN_00312) was located upstream of the resolvase gene (MNHDEDAN_00313) in a divergent orientation, potentially completing the canonical Tn3 structure. The corresponding DUF536 protein was, however, truncated in 4S77, suggesting the element there is likely non-functional.

#### Genomic islands (GIs)

Three GIs were identified on the chromosome of 4S77 and two on 4S90 (**Fig 8A**), encoding genes involved in transport, cell wall modification, and mobile genetic elements, along with several hypothetical proteins.

**Fig 8.**
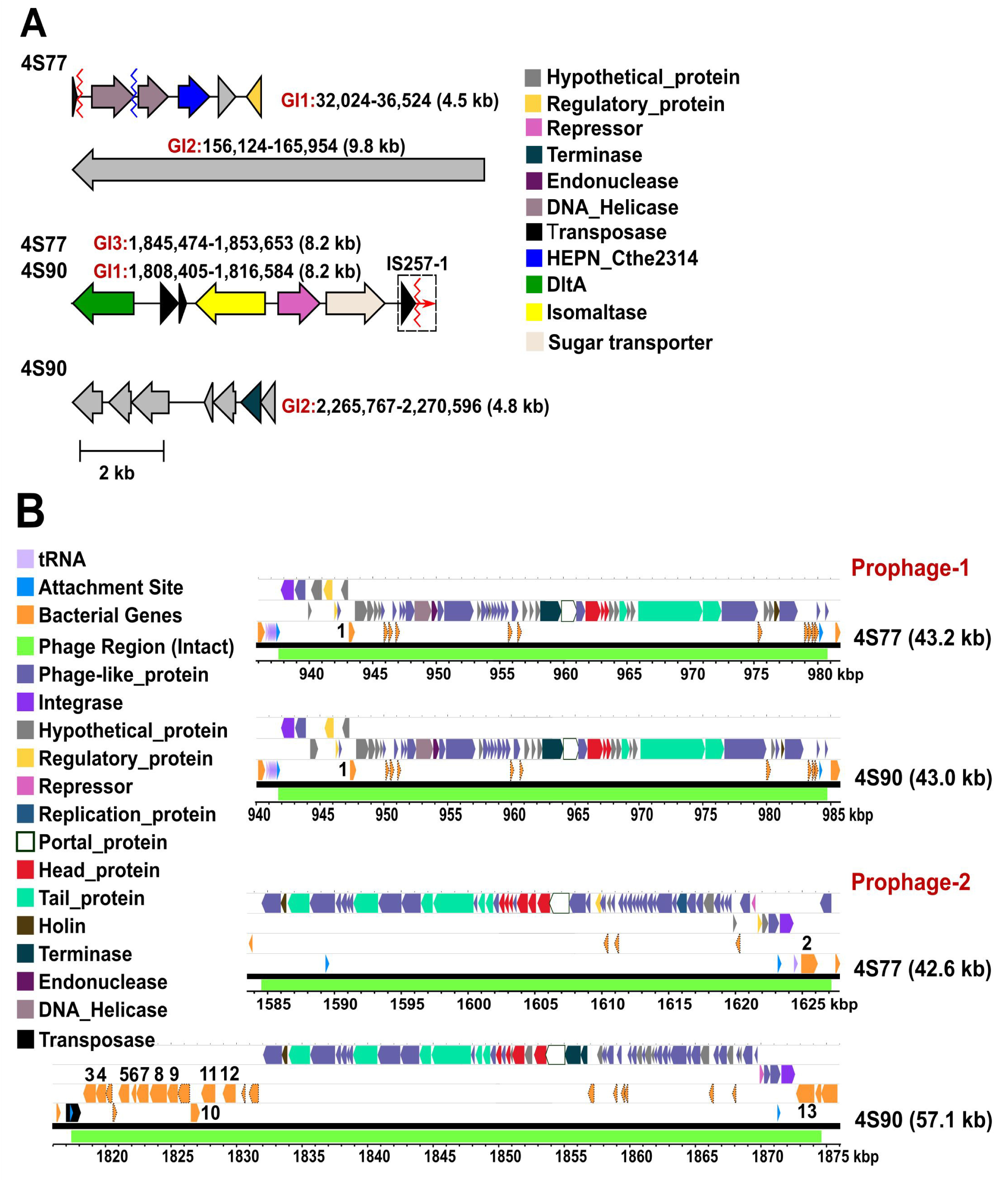
Genomic Islands and Prophage Regions in *Staphylococcus chromogenes* Strains 4S77 and 4S90. (A) Genomic Islands (GIs) of *Staphylococcus chromogenes* strains 4S77 and 4S90. Dlt, D-alanyl transfer protein; HEPN_Cthe2314, Cthe_2314 HEPN (higher eukaryotes and prokaryotes nucleotide-binding) domain-containing protein; MFS, major facilitator superfamily; IS257-1, the IS-6 family insertion sequence downstream of 4S90-GI1. Red and blue zigzag lines indicate truncated and fragmented genes, respectively. “Truncated” indicates predicted transposase genes with incomplete open reading frames (ORFs). “Fragmented” indicates transposase sequences split into two separate ORFs due to internal disruptions; (B) Prophage regions of *S. chromogenes* strains 4S77 and 4S90. Protein-coding bacterial genes are numbered as follows: 1. Endonuclease; 2. Aminoacyltransferase; 3. Dehydrogenase; 4. Hydrolase (YutF); 5. Insertase (YidC); 6. antitoxin (YutD); 7. Lipoyl synthase; 8. Bifunctional metallophosphatase/5’-nucleotidase; 9. Sulfite exporter (TauE/SafE); 10. Hydrolase; 11. Nitronate monooxygenase; 12. Hemolysin III family protein, 13. Iron-sulfur cluster assembly protein (SufB).

In strain 4S77, GI1 spanned approximately 4.5 kb and encoded five open reading frames (ORFs). The final three genes encoded a HEPN (higher eukaryotes and prokaryotes nucleotide-binding) domain-containing protein, a hypothetical protein, and a predicted regulatory protein. Comparative analysis with ten publicly available *S. chromogenes* genomes revealed that five strains (76ME, CCM, DSM, IM, and IVB6199) harbored a region corresponding specifically to these last three ORFs, suggesting partial conservation of this sub-region. Notably, the HEPN-domain protein and adjacent hypothetical protein were reminiscent of the HEPN-transmembrane (TM) antiphage defense system described in *S. saprophyticus* strain SS413 [30]. However, neither the HEPN protein (245 aa) nor the hypothetical protein (136 aa) in 4S77 showed significant sequence similarity to their *S. saprophyticus* counterparts (WP_129531117.1 and WP_129531118.1, respectively), and the hypothetical protein lacked a predicted TM domain. While homology appeared unlikely, the possibility of an analogous antiphage role cannot be excluded and may merit further investigation.

In strain 4S77, GI2 spanned approximately 9.8 kb and encoded a single ORF annotated as a hypothetical protein. Homologous nucleotide sequences were identified in nine of the ten publicly available *S. chromogenes* genomes (absent only in IVB6199). In these strains, the region was frequently annotated as encoding an LPXTG cell wall anchor domain-containing protein. However, in all cases, the gene appeared fragmented, disrupted by frameshifts or premature stop codons, resulting in inconsistent ORFs and low aa-level conservation. This suggests that the region represents a decaying cell wall-associated locus that has undergone independent pseudogenization events across the species.

GI3 in strain 4S77 and GI1 in strain 4S90 were syntenic genomic islands comprising eight ORFs. In addition to *dltA*, which is part of the conserved *dltABCD* operon found in all *S. chromogenes* strains, this island included three accessory genes encoding an isomaltase, a transcriptional repressor, and a sugar transporter. These three genes were present only in strains BTM, IM, and IVB6199; the remaining seven genomes lacked them entirely, indicating a discrete loss of this accessory module. Notably, the island was flanked by a truncated IS6 family transposase upstream and an intact IS1182 family transposase downstream, an arrangement consistent with the chromosomal IS configuration previously described and suggestive of potential mobilization of this region through an IS-mediated or hitchhiking mechanism.

In strain 4S90, GI2 spanned approximately 4.8 kb (positions 2,265,767–2,270,596 bp) and encoded seven ORFs, six of which were annotated as hypothetical proteins. The second-to-last ORF encoded a predicted terminase. Comparative analysis with ten *S. chromogenes* genomes revealed a heterogeneous presence–absence pattern, with different subsets of genes detected in several strains, including 17A, BTM, CCM, IVB6199, and IVB6200, but no complete copy was identified.

Despite the presence of accessory genes with predicted roles in transport, metabolism, and mobile element activity, none of the identified genomic islands in either strain encoded known virulence determinants, nor did they exhibit features typically associated with pathogenicity.

#### Prophage regions

PHASTEST analysis identified two intact chromosomal prophage regions in each of the *S. chromogenes* strains 4S77 and 4S90 (**Fig 8B, S11 Table**). These prophages ranged from 42.6 to 57.1 kb in length and carried typical phage-associated modules, including integrase, terminase, capsid, tail, and lysis genes, along with a variable number of bacterial genes. Prophage region 1 was located at similar chromosomal positions in both strains and spanned 43.2 kb in 4S77 and 43.0 kb in 4S90. The two regions exhibited 85% coverage and 97% nucleotide identity, with broadly conserved gene content, suggesting they likely originated from a common ancestral phage. In both strains, the *attL* site was located within a tRNA^Ser^ gene and the *attR* site appeared as a direct repeat in an intergenic region, consistent with site-specific integration. Each prophage contained over 50 predicted phage-related coding sequences, including structural and replication components. In 4S77, 11 bacterial CDSs were identified, while 4S90 contained ten. Of these, only one gene, annotated as an endonuclease (bacterial gene 1), had a defined function; the rest were annotated as hypothetical proteins.

The second prophage regions in strains 4S77 and 4S90 measured 42.6 kb and 57.1 kb, respectively. BLASTn alignment between the two revealed 44% coverage with 94% nucleotide identity, restricted to the phage-related portions, indicating strong conservation of the core phage backbone. Outside this shared segment, the regions differed in bacterial gene content: 4S77 encoded four bacterial coding sequences, of which only one was functionally annotated as aminoacyltransferase, while 4S90 contained 22 bacterial CDSs, 11 of which had assigned functions based on database annotations.

Despite the presence of several bacterial genes within the prophage regions of both strains, no canonical virulence factors were identified. In particular, a gene encoding a hemolysin III (Hly-III) family protein was found in the second prophage region of strain 4S90. Although annotated as a member of the Hly-III family, TM topology predictions indicated a structure comprising a signal peptide, three TM helices, and a large cytoplasmic C-terminal domain (**S4A Fig**). This topology was distinct from that of the Hly-III protein from *Bacillus cereus* (UniProt: P54176), which lacks a signal peptide and contains seven TM helices (**S4B Fig**), consistent with its proposed pore-forming activity [31]. However, even in *B. cereus*, secretion of hemolysin III has not been experimentally demonstrated, and its contribution to virulence remains unproven and speculative [32]. Furthermore, BLASTp analysis showed no significant sequence similarity between the two proteins, suggesting that the 4S90-encoded Hly-III-like protein likely represents a functionally distinct, membrane-associated paralog without direct hemolytic or pathogenic activity.

#### Staphylococcal cassette chromosome *mec* (SCC*mec*)

No SCC*mec* element was detected in either strain using SCC*mec*Finder. The same result was obtained for all ten publicly available *S. chromogenes* genomes, which were likewise predicted to be methicillin-susceptible. In all cases, SCC*mec*Finder reported the absence of both *mecA* and *mecC*, consistent with the lack of this resistance element across the species dataset.

#### Plasmids

In strains 4S77 and 4S90, the shared ∼57 kb plasmid encoded a lipase and two adjacent operons comprising the ribulose monophosphate (RuMP) pathway and the oxidative branch of the pentose phosphate pathway (**S12 Table**). These operons appeared functionally interconnected: the RuMP pathway fixes formaldehyde into fructose 6-phosphate [33], while the pentose phosphate pathway generates ribulose 5-phosphate and NADPH. Although chromosomal copies of the RuMP pathway were also present in 4S77, 4S90, and all included *S. chromogenes* genomes, the plasmid-encoded versions may reflect gene duplication or enhanced expression under specific environmental conditions. Notably, the operon encoding the oxidative pentose phosphate pathway lacked 6-phosphogluconolactonase, and metabolic pathway analysis using KEGG confirmed that the gene was also absent from the chromosome. A plasmid-borne α/β-hydrolase fold esterase may substitute this function, either through promiscuous catalytic activity or structural flexibility. Alternatively, lactone hydrolysis may proceed non-enzymatically, albeit inefficiently [33, 34]. In strain 4S90, a smaller plasmid additionally harbored a lincosamide resistance determinant. Across the broader set of *S. chromogenes* genomes, plasmids varied in replicon type and gene content, but functional genes were infrequent and scattered. Lipase was the only consistently plasmid-encoded virulence-associated function, while resistance determinants were rare and strain-specific. No consistent association was observed between replicon type and the presence of resistance or virulence-related functions (**S13 Table**). These findings indicate that plasmids play a minimal and sporadic role in mediating antimicrobial resistance and virulence potential in *S. chromogenes*.

### Bacteriocin biosynthetic gene clusters

Mining the genomes of *S. chromogenes* strains 4S77 and 4S90 revealed two putative BGCs encoding ribosomally synthesized and post-translationally modified peptides (RiPPs) on the 57-kb plasmid, as predicted by antiSMASH 5.0. The first RiPP-like cluster was 12.1 kb in size and showed limited similarity to the *lactocin S* gene cluster from *Lactobacillus sakei* (16 kb). The similarity was restricted to transposase-related regions, with no homology to the structural gene *lasA*. The cluster included two small ORFs encoding peptides of 42 and 51 amino acids, located adjacent to an ABC transporter with a 150-residue N-terminal extension corresponding to a peptidase domain (**S5A Fig**).

Both peptides contained a 15-residue N-terminal leader of the double-glycine type (XXXLSXXELXXIXGG), differing only at position 11 (serine replaced by asparagine), and helix–helix-interacting GxxxG motifs (**S5B Fig**). These are features characteristic of class IIb, two-peptide bacteriocins, in which the adjacent ABC transporter cleaves the leader at the GG motif and simultaneously translocates the mature peptides across the cell membrane features that are characteristic of class IIb, the two-peptide, bacteriocins [35]. While these sequence features supported a potential export and processing mechanism, signal peptide predictions were inconsistent: the 42-amino acid (aa) peptide showed a cleavage site at the GG motif, whereas the 51-aa peptide lacked a predicted signal sequence. BLASTp analysis revealed low similarity (29% identity) between the 42-aa peptide and peptide A (31 aa) from aureocin A70, a four-component class II bacteriocin produced by *S. aureus* (**S5C Fig**). Notably, the *a70* operon encodes an ABC transporter (*aurT*) lacking peptidase activity, consistent with the absence of leader sequences in aureocin A70 peptides [36].

The second BGC identified in *S. chromogenes* strains 4S77 and 4S90 was associated with the production of the thiopeptide micrococcin P1 (MP1). AntiSMASH analysis and manual annotation revealed a conserved gene cluster spanning approximately 11 kb and comprising 12 genes involved in MP1 biosynthesis, located on the 57-kb plasmid. The cluster exhibited high synteny and protein homology with previously characterized MP1 gene clusters from *S. hominis* S34-1 [37], *S. equorum* KAVA [38], *S. agnetis* 4244 [39], and *Macrococcus caseolyticus* 115 [40], and showed partial conservation with the more complex, chromosomally encoded BGC of *B. cereus* ATCC 14579 [41] (**Fig 9**). Gene nomenclature was assigned based on the original *tcl* gene cluster described in thiocillin-producing strains, following established conventions [42]. The precursor peptide was encoded by *tclE* and comprised a 35-residue leader sequence at the N-terminus, followed by a highly conserved 14-residue core peptide, shared across MP1-producing species, which undergoes extensive post-translational modification (**Fig 10AB**). The genes *tclS* and *tclP* encode dehydrogenases involved in the modification of the C-terminal threonine residue of the precursor peptide during maturation. The cluster also comprised *tclI*, *tclJ*, and *tclN*, which together catalyse the conversion of cysteine residues into thiazole rings. TclI is predicted to contain a RiPP recognition element, enabling binding to the leader sequence of the precursor peptide and thereby assisting in recruiting the modification enzymes. TclJ mediates the ATP-dependent formation of thiazoline intermediates, which are subsequently oxidised by TclN to yield thiazole heterocycles. *tclK* and *tclL* are predicted to encode dehydratases responsible for converting serine and threonine residues into dehydroalanine and dehydrobutyrine. TclM is likely responsible for catalysing the cyclisation that generates the central pyridine ring and initiates leader peptide cleavage. Regulatory control may be provided by *tclU*, which encodes a putative MerR-family transcriptional regulator with a helix-turn-helix motif. Self-immunity is presumed to be conferred by *tclQ*, a homologue of ribosomal protein L11 [42]. The sequence alignment of L11 and TclQ proteins (**S6 Fig**) provided evidence for the proposed self-resistance mechanism. TclQ is a paralogue of the ribosomal protein L11, encoded within the MP1 biosynthetic gene cluster. Unlike the native L11, TclQ carries a conserved proline-to-threonine substitution within the N-terminal domain, a region critical for binding MP1. MP1 targets the GTPase-associated centre of the ribosome, binding between L11 and the 23S rRNA, where it interferes with elongation factor G and blocks translation. The substitution in TclQ is predicted to reduce MP1 binding, thereby rendering the ribosome less sensitive to its inhibitory effects [42]. This suggests that self-immunity in *S. chromogenes* is achieved through the expression of TclQ, which likely replaces the native L11 in the ribosome, shielding the producing strain from its own antimicrobial compound.

**Fig 9.**
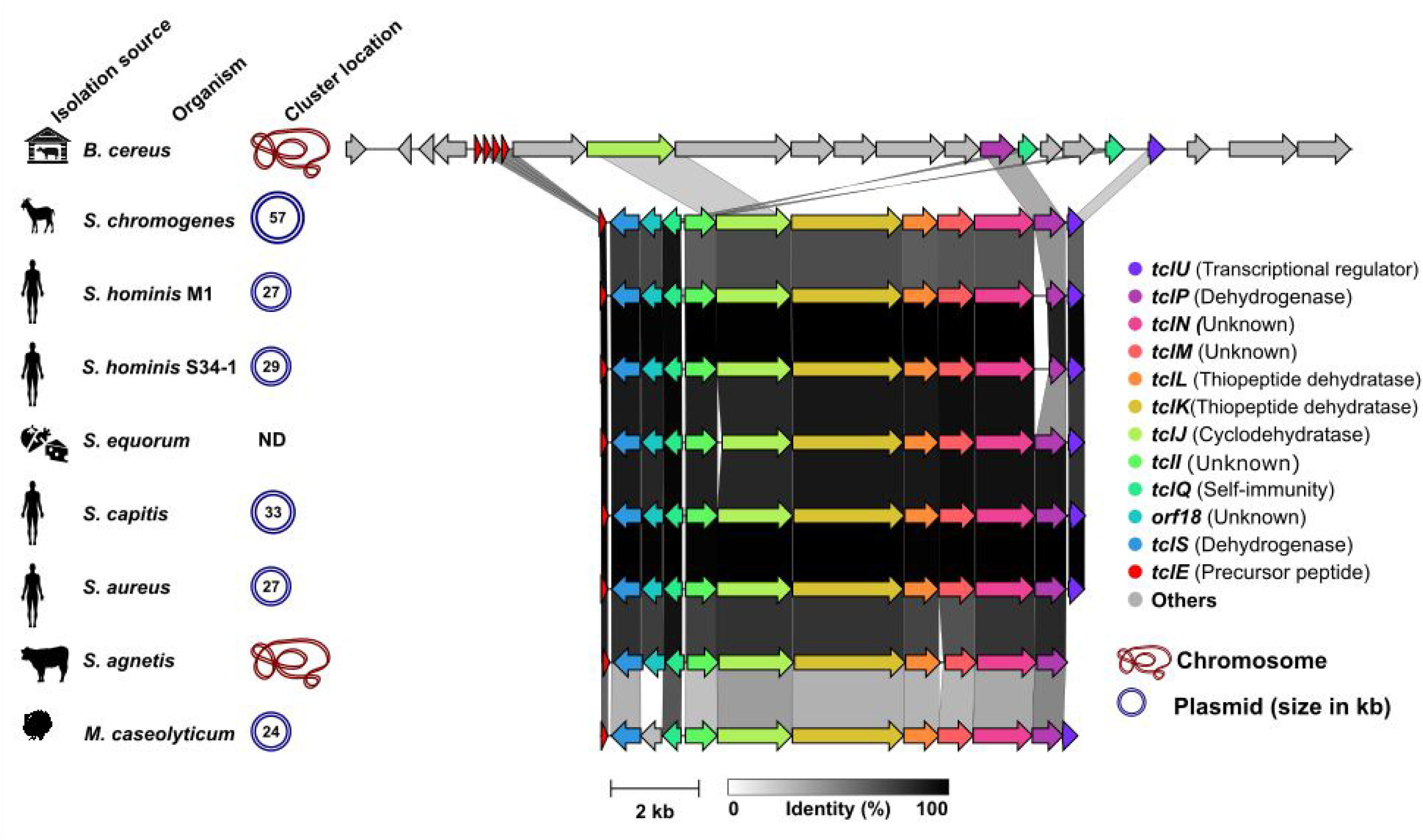
Comparative analysis of micrococcin P1 biosynthetic gene clusters across diverse bacterial species. The gene cluster identified in *Staphylococcus chromogenes* strains 4S77 and 4S90 is shown alongside homologous clusters from *Bacillus cereus* ATCC 14579 (GenBank accession no. CP034551), *S. hominis* M1 (this study), *S. hominis* S34-1 (CP040733), *S. equorum* KAVA (MW633237), *S. capitis* Sc1516939 (CP145210), *S. aureus* UP_1591 (CP047810), *S. agnetis* 4244 (JABULG020000000), and *Macrococcus caseolyticus* 115 (KM613043). The sequence from *Mammaliicoccus sciuri* IMDO-S72 was not available and could not be included. Clusters were identified on either plasmids or chromosomes. Gene functions are color-coded as indicated in the legend. The core biosynthetic genes (*tclE*–*tclQ*) exhibit strong conservation across *Staphylococcus* species, with sequence similarity illustrated by shaded grey connectors reflecting amino acid identity. ND, not determined.

**Fig 10.**
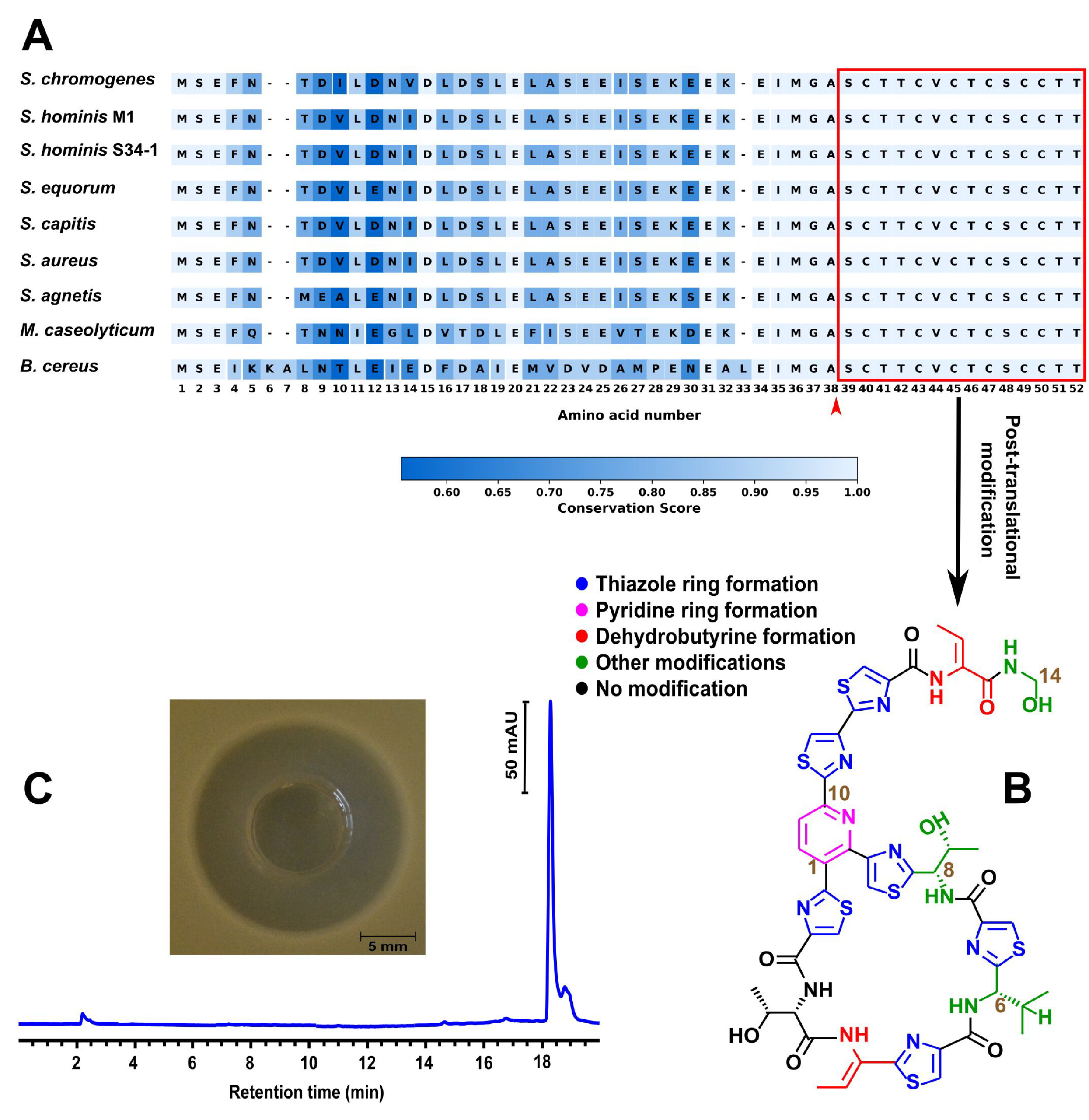
Sequence conservation, structure, and functional validation of micrococcin P1 (MP1) from *S. chromogenes* strains 4S77 and 4S90. (A) Alignment of the precursor peptide encoded by *tclE* from the MP1 cluster in *S. chromogenes* strains 4S77 and 4S90 with homologous sequences from *Bacillus cereus* ATCC 14579 (CP034551), *S. hominis* M1 (this study), *S. hominis* S34-1 (CP040733), *S. equorum* KAVA (MW633237), *S. capitis* Sc1516939 (CP145210), *S. aureus* UP_1591 (CP047810), *S. agnetis* 4244 (JABULG020000000), and *Macrococcus caseolyticus* 115 (KM613043). The sequence from *Mammaliicoccus sciuri* IMDO-S72 was not available and could not be included. The red arrow marks the predicted cleavage site between the leader and core regions, with the red box indicating the conserved 14-residue core peptide subject to post-translational modification; (B) Schematic representation of post-translational modifications on the 14-residue core peptide leading to mature MP1. Colour legend indicates the different modification types. Adapted from Wieland Brown *et al.* [41]; (C) RP-HPLC chromatogram of the purified compound from *S. chromogenes* strains 4S77 and 4S90, showing a dominant peak at the expected retention time for MP1. Inset shows the inhibition zone produced by this purified sample (2.5 µg) against *S. aureus* JE2.

In addition to the two plasmid-encoded RiPP clusters, a subtilosin A-like operon was identified on the chromosome of *S. chromogenes* 4S77, spanning base positions 235,620 to 254,823. BAGEL4 analysis revealed partial similarity to the subtilosin A biosynthetic locus from *B. subtilis* 168. However, only two genes were detected within the region: a predicted structural gene encoding the precursor peptide (*sboA*) and a putative immunity gene (*albB*) (**S7A Fig**). Alignment of the *S. chromogenes* cluster with the full subtilosin operon in *B. subtilis* showed that it lacks the remaining *alb* genes (*albA–albG*) required for post-translational modification and peptide maturation. These include enzymes that are responsible for cyclisation and the formation of unique sulfur-to-α-carbon thioether bonds, which are essential for biological activity [43]. The precursor peptide shared limited similarity with the subtilosin A sequence from *B. subtilis* (**S7B Fig**), and BLASTp analysis of the putative immunity protein against AlbB revealed only 43.5% identity over 80% coverage, with a weak E-value (6e–08) and low bit score (29.6). Taken together, these findings suggest that the cluster in *S. chromogenes* 4S77 represents an incomplete or degenerate operon, possibly acquired through horizontal gene transfer, and is unlikely to support the biosynthesis of an active subtilosin-like compound.

### Analytical confirmation of MP1 as the antibacterial compound in *S. chromogenes*

Of the three identified RiPP-like clusters in *S. chromogenes* strains 4S77 and 4S90, only the MP1 BGC was linked to the production of an active antimicrobial compound. Culture supernatants (SN) from *S. chromogenes* strains 4S77 and 4S90 were extracted using ethyl acetate. Antimicrobial testing revealed that only the organic phase retained activity. The extracts were subsequently purified and reversed-phase high-performance liquid chromatography (RP-HPLC) analysis revealed a dominant peak matching that of the MP1 standard. This purified preparation also displayed antimicrobial activity against *S. aureus* USA300 JE2 (**Fig 10C**). liquid chromatography–mass spectrometry (LC-MS) analysis of the purified compound revealed a prominent ion at mass-to-charge ratio (*m/z*) 572.6, corresponding to the doubly charged [MP1 + 2H]²⁺ ion, and a secondary ion at *m/z* 1144.2, corresponding to the singly charged [MP1 + H]⁺ ion (**S8 Fig**), consistent with the expected molecular mass of MP1 [40].

By contrast, synthetic peptides from the unidentified RiPP-like cluster, with and without leader sequences, were tested individually and in equimolar combinations against *Micrococcus luteus* but showed no detectable activity. In comparison, the aureocin A70 peptide A, used as a positive control, exhibited strongly inhibitory activity.

Subtilosin A, being highly hydrophobic and nearly insoluble in aqueous solvents [44], would be expected to partition into the ethyl acetate phase during extraction; however, no corresponding compound was detected, supporting that the predicted cluster is non-functional due to the absence of essential biosynthetic genes.

### Distribution and uniqueness of MP1 gene cluster in *Staphylococcus*

To explore the distribution of MP1 BGC across the *Staphylococcus* genus, 2,939 complete genomes were screened using a tBLASTn search with the MP1 precursor peptide sequence from *S. chromogenes* 4S77 and 4S90. The dataset included isolates from diverse sources, with 69% derived from human-associated samples. Notably, 2,175 of the genomes belonged to *S. aureus* (74%), reflecting its dominance in public databases (**Fig 11A, S14 Table**). This approach identified nine genomes with significant hits (0.3%). All nine were human-derived and included one *S. hominis*, one *S. capitis*, and seven *S. aureus* strains. Further analysis confirmed that each of these strains harbored a complete MP1 cluster, with conserved gene content and synteny. In all cases, the clusters were plasmid-encoded, with plasmid sizes ranging from 13 to 33 kb (**Fig 11B**). While these findings suggest that MP1-like BGCs are rare within *Staphylococcus*, their apparent restriction to human-associated strains, particularly *S. aureus*, may, in part, reflect the taxonomic and sampling bias of the available genomic dataset. To further assess whether the MP1 cluster is unique to *S. chromogenes* 4S77 and 4S90, a BLASTp search was conducted against the NCBI non-redundant protein database, restricted to the taxon *S. chromogenes*. No additional hits were detected, indicating that this cluster has not been recorded in any other *S. chromogenes* isolate in the available NCBI protein database.

**Fig 11.**
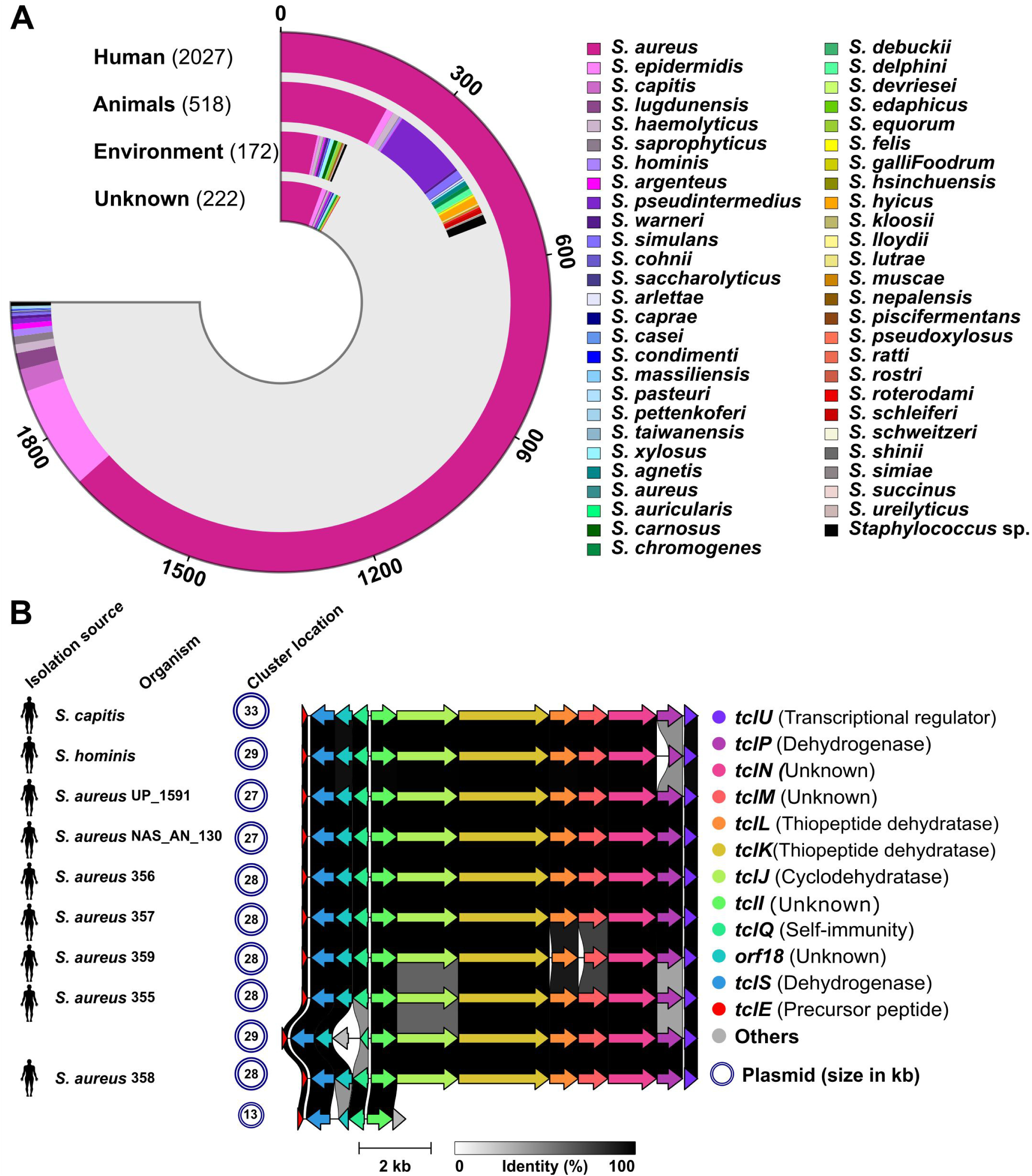
Distribution and organisation of micrococcin P1 (MP1) biosynthetic gene clusters in *Staphylococcus* genomes. (A) Source distribution and species breakdown of 2,939 *Staphylococcus* genomes with complete assemblies retrieved from the NCBI database; (B) Comparative gene cluster analysis of the nine genomes identified to encode a complete MP1 biosynthetic gene cluster based on tBLASTn screening with the precursor peptide from *S. chromogenes* strains 4S77 and 4S90. All clusters were plasmid-encoded and exhibited conserved gene content and synteny. Arrows represent individual genes, colour-coded by predicted function as shown in the legend. Sequence similarity illustrated by shaded grey connectors reflecting amino acid identity.

## Discussion

Antimicrobial activity in *S. chromogenes* was first reported in isolates capable of inhibiting *S. aureus*, *Streptococcus uberis*, and *Streptococcus dysgalactiae*, indicating the presence of diffusible inhibitory compounds [13]. Cytolysin-like and nukacin-like bacteriocins have since been identified in *S. chromogenes*, though their genetic basis and biological function remain uncharacterized [1, 11]. More recently, the type strain ATCC43764 was shown to produce 6-thioguanine, a purine analogue that impairs *S. aureus* virulence through inhibition of quorum sensing and purine biosynthesis [12]. In this context, the identification of a complete, plasmid-encoded MP1 BGC in two goat milk isolates provides direct evidence of a distinct antimicrobial mechanism in *S. chromogenes*, albeit one that appears to be extremely rare within the genus (0.3% of complete *Staphylococcus* genomes) and absent from all other *S. chromogenes* examined. This low frequency is consistent with previous large-scale screening study, in which MP1 production was detected in only four out of 890 *Staphylococcus*/*Mammaliicoccus* isolates (0.4%) from diverse origins and species [45]. MP1 BGCs have been reported in several *Staphylococcus* species, including *S. agnetis* [39], *S. hominis* [37, 45], *S. aureus* [45], and *S. equorum* [38, 46], as well as in the closely related *Macrococcus caseolyticus* [40] and the reclassified *Mammaliicoccus sciuri* [42]. All characterized clusters in these taxa share a highly conserved core structure and gene arrangement with that found in *S. chromogenes*. In a more complex arrangement, the chromosomal MP1 cluster of *B. cereus* ATCC 14579 spans 22 kb and includes 24 genes, among them four precursor peptide genes (*tclE, tclF, tclG,* and *tclH*), two L11-type immunity genes (*tclQ* and *tclT*), and several additional modification enzymes (*tclD, tclO,* and *tclV*) not found in staphylococcal-type clusters [41]. These features may underlie the production of eight structurally distinct thiopeptides by *B. cereus*, in contrast to the single MP1 compound encoded by the more streamlined staphylococcal BGCs.

*In vitro*, MP1 has shown potent activity against a broad range of Gram-positive bacteria, including clinically relevant *Staphylococcus*, *Streptococcus*, *Enterococcus*, and *Listeria* species, as well as skin-associated opportunists and food-related lactic acid bacteria [17, 37, 38, 47]. Activity has also been demonstrated against *Mycobacterium tuberculosis* [48], but not against Gram-negative species. *In vivo*, MP1 has also shown efficacy in murine models of methicillin-resistant *S. aureus* (MRSA) skin infection, administered as a purified compound, via an MP1-producing strain, or in combination with rifampicin [37, 38].

Given its broad antimicrobial activity and demonstrated efficacy in infection models, a key question is how MP1 production may function in the natural ecology of *S. chromogenes*. Unlike other MP1-producing staphylococci, *S. chromogenes* is primarily associated with the ruminant udder, where it is consistently recovered from milk and teat skin, yet rarely isolated from non-host environments [6]. Human isolation of *S. chromogenes* is exceedingly rare [49], with no reported infections to date. Among 177 genomes with metadata in NCBI, only one was of human origin, isolated from a pet café worker. Reflecting this host-restricted distribution, the species is classified as Risk Group 1 in several countries, while Germany assigns it to Risk Group 2 due to potential occupational exposure risks [50]. Its persistent presence in the udder environment suggests a species well adapted to the ruminant mammary niche. This colonization capacity may be supported by biofilm formation. All examined *S. chromogenes* genomes encoded the autolysin Atl, a known initiator of biofilm formation, as well as FnB, Bap, and sortase A, the latter anchoring a wide range of LPXTG-motif surface proteins, including FnB and Bap, to the cell wall [51, 52]. In addition to these core components, all 12 complete genomes also encoded enolase, a glycolytic enzyme with a secondary, moonlighting role in biofilm formation through surface association and stabilization of the extracellular matrix [53]. Given the apparent absence of the *ica* operon and the accumulation-associated protein Aap, biofilm formation in *S. chromogenes* likely relies on the combined action of Atl, FnB, Bap, sortase A, and enolase. This multifactorial system may underpin the observed variability in biofilm phenotypes, which range from absent to strong among *S. chromogenes* strains [9, 54]. All genomes also encoded the PGA capsule, a surface polymer reported to protect against antimicrobial peptides and neutrophil phagocytosis, suggesting this feature may also contribute to persistence within the host [55].

*S. chromogenes* also exhibits a versatile and resilient metabolism, shaped for survival in the nutrient-limited, microbially competitive environment of the ruminant udder. Genomic data revealed that this metabolic resilience is supported by enriched KOs and expanded amino acid biosynthesis pathways, aligned with the challenges of surviving in the nutrient-poor environment of healthy udder. Unlike mastitic udder, healthy udder contains very low levels of free amino acids, particularly BCAAs, AAAs, and lysine [56]. *S. chromogenes* likely offsets this scarcity through de novo synthesis, using the *ilv-leu* operon (BCAAs), the shikimate-AAA pathway (AAAs), and *dapABD/lysA* (lysine). Histidine, one of the few amino acids more available in healthy udder, may serve as an alternative nitrogen and carbon source via the hut operon. The presence of the histidine biosynthesis operon (*his*), along with various amino acid permeases, on the other hand, suggests a dual strategy of biosynthesis and nutrient scavenging. Enrichment in the pantothenate-CoA biosynthesis pathway supports metabolic integration, with BCAAs feeding directly into branched-chain fatty acids (BCFA) production. CoA, derived from pantothenate, is critical for BCFA production and broader stress responses. These BCFAs are key to maintaining membrane fluidity, integrity, and function, especially under environmental stress. BCAAs also regulate the global nutrient sensor CodY, with BCAA shortages impacting protein production, membrane lipid synthesis, and overall fitness [57]. Additional enrichment in lysine and D-amino acid metabolism, both involved in cell wall cross-linking and remodeling, points to mechanisms for structural adaptation under stress [58, 59]. Together, these pathways highlight how metabolic flexibility supports not only nutrient acquisition but also cell envelope robustness, critical traits for persistence in the udder niche.

Collectively, these colonization-associated traits reflect a multifaceted survival strategy fine-tuned to the udder environment. Beyond structural defenses, functional mechanisms such as bacteriocin production may further enhance niche persistence by conferring a competitive advantage against mastitis-associated bacteria [1, 11, 60]. Evidence for such a role is provided by studies on *S. aureus*, where horizontal acquisition of the aureocin A70 locus has been linked to increased fitness during mammary gland colonization [61]. Furthermore, *S. chromogenes* has been shown to possess enhanced ecological competitiveness through the secretion of non-microbicidal compounds that inhibit pathogen biofilm formation, a hallmark commensal strategy for protecting epithelial niches, regardless of its own biofilm-forming capacity [9, 60]. Beyond biofilm interference, *S. chromogenes* exoproducts have been shown to downregulate key *S. aureus* virulence genes by disrupting the *agr* quorum sensing system and attenuating of pathogenic potential [62]. One such exoproduct is the autoinducing peptide with a core region of “SINPCTGFFC”, recently shown to potently inhibit all *S. aureus* and *S. epidermidis agr* variants [63]. In addition to this reported variant, other sequences were identified in the genomes studied here, including “AMDPCTGFF” and “SINPCTAFF”; notably, strains 4S77 and 4S90 carried the former. As the conserved core motif is retained, these variants may remain active, though this requires experimental confirmation. Recent findings further suggest that certain *S. chromogenes* strains can also reduce *S. aureus* internalization into mammary epithelial cells through mechanisms that appear independent of *agr* regulation [64]. These antagonistic effects have also been observed *in vivo*, where *S. chromogenes* exoproducts reduced *S. aureus* colonization in a murine mastitis model [2].

Moreover, *S. chromogenes* has been shown to inhibit colonization by mastitis pathogens not only through direct antagonism but also by modulating the host innate immune response, an effect that may simultaneously hinder pathogen establishment and promote its own persistence within the udder environment [4, 8, 65]. While capable of activating core neutrophil effector functions, phagocytosis, reactive oxygen species production, bacterial killing, and release of neutrophil extracellular traps, *S. chromogenes* elicited a comparatively attenuated inflammatory response compared to major mastitis pathogens. Neutrophil recruitment was markedly reduced, granule enzyme release remained limited, and pro-inflammatory cytokines, particularly interleukin-6, were comparatively suppressed. This restrained activation may allow *S. chromogenes* to persist as a discreet colonizer, evading clearance while maintaining a foothold in the udder niche [7, 66]. This relatively subdued immunological profile is consistent with our genomic findings, which revealed that all examined *S. chromogenes* strains lacked the majority of known staphylococcal virulence genes, particularly those associated with aggressive pathogenicity. Even among the limited set of virulence genes detected, phenotypic assays showed that nuclease, hemolytic, and lipase activities were consistently weaker than those of *S. aureus*. While this reduced activity under controlled conditions supports the view of *S. chromogenes* as a less aggressive species, both transcriptional activation and phenotypic expression are shaped by host-specific cues, nutritional status, microbial interactions, and other environmental factors, any of which could modulate virulence expression *in vivo* [51]. The functional contribution of these virulent traits to the biology of *S. chromogenes*, particularly in the context of host interactions, has yet to be clearly defined.

Whether the ecological persistence of *S. chromogenes* reflects a trait of a mastitis pathogen or a commensal strategy that shields the host from more aggressive invaders remains unresolved. One explanation may lie in its strain-level diversity: *S. chromogenes* isolates have been shown to differ in their ability to modulate innate immunity, inhibit pathogen biofilms, and interact with mammary epithelial cells [5, 7–10]. These functional differences point to a species that, depending on the strain, can act as either a microbial bodyguard or a low-key saboteur. Much like a biological gatekeeper, *S. chromogenes* may occupy space, calm host defenses, and quietly hold the line against more virulent foes, but the balance it strikes is still an open question. Clarifying this ambiguity will require strain-specific studies in biologically relevant models that reflect the complexities of the mammary gland.

## Conclusion

This study characterized two goat-derived *S. chromogenes* strains, 4S77 and 4S90, that produce the thiopeptide bacteriocin MP1, a feature previously undocumented in this species. Genomic and phenotypic analyses showed that both strains pair potent antimicrobial activity with low virulence, lacking classical pathogenicity markers and exhibiting susceptibility to antibiotics. Their conserved metabolic capacity, colonization-associated genes, and production of MP1 point to a dual ecological role: persistent commensals within the mammary gland and potential microbial gatekeepers against mastitis pathogens. These findings support the idea that some *S. chromogenes* strains may contribute to maintaining microbial balance and udder health in dairy ruminants.

## Methods and materials

### Genomic DNA isolation

To prepare genomic DNA, overnight bacterial cultures were first subjected to enzymatic lysis using lysozyme (2.5 mg/mL) and lysostaphin (1.25 mg/mL) at 37°C for 30 minutes. DNA extraction was then performed with the Quick-DNA™ Microprep Kit (ZYMO Research, Freiburg, Germany), following the manufacturer’s instructions. DNA concentration and purity were assessed using a NanoPhotometer® NP80 spectrophotometer (Implen, Munich, Germany).

### Whole genome sequencing, de novo assembly, and annotation

Whole genome sequencing of the *S. chromogenes* strains 4S77 and 4S90 was performed by the NGS Competence Center Tübingen (NCCT), University of Tübingen, Germany. Genomic DNA was prepared for sequencing using the Illumina Nextera DNA Flex kit (short reads) and the Oxford Nanopore Technologies (ONT) Native Ligation kit (long reads). Sequencing was carried out on an Illumina MiSeq system with the MiSeq® Reagent Kit v2 (300 cycles) and on an ONT MinION device using MinION Flow Cells.

A hybrid assembly was conducted with Unicycler (v0.4.4) [67], integrating Illumina paired-end reads and ONT long reads. The pipeline used SPAdes (v3.13.2) [68] for short-read assembly, miniasm [69] for long-read scaffolding, and Racon [70] for iterative polishing. Both strains yielded complete, circularized genome assemblies. Final assemblies were further refined to remove repetitive sequences at contig boundaries.

To complete the genome analysis, structural plasmid elements were confirmed using plasmIDent [71], which aligned full-length Nanopore reads to support the presence of plasmid sequences. Annotation of the finalized assemblies was then performed using Prokka version 1.13 [72]. Circular genome maps for *S. chromogenes* strains 4S77 and 4S90 were generated using the Python package pyCirclize tool [73] was used to create circos-style plots of annotated genomic features following hybrid assembly and annotation.

### Phylogenetic Analysis

Phylogenetic relationships among *S. chromogenes* and other *Staphylococcus* species were inferred using the Codon Tree pipeline available in PATRIC [74]. The analysis was performed using 100 randomly selected single-copy genes from PATRIC’s global protein families (PGFams), shared across all selected genomes. Protein sequences were aligned using MUSCLE, and corresponding nucleotide sequences were aligned with the codon-aware Codon_align function from BioPython. A concatenated alignment of amino acid and nucleotide sequences was used to construct a maximum-likelihood tree in RAxML, using a partitioned model and 100 rounds of rapid bootstrapping for branch support. The analysis included 56 complete genomes, comprising 44 reference *Staphylococcus* species, ten publicly available *S. chromogenes* genomes, and the two sequenced isolates 4S77 and 4S90. To investigate relationships specifically within *S. chromogenes*, a core-genome phylogeny was constructed using 177 publicly available genomes compiled from NCBI (retrieved on 16 May 2025) alongside the two isolates from this study. The core genome was identified and aligned using Parsnp from the Harvest software package (v1.7.4) [75]. High-confidence core-genome SNPs identified by Parsnp were then used to reconstruct the phylogeny using FastTree 2 [76]. The resulting trees were visualized with the Interactive Tree of Life (iTOL) [77].

### Metabolic pathway analysis

To compare the metabolic capabilities of *Staphylococcus* strains, KO annotations were generated for each genome using Bakta [78], ensuring consistent annotation across all groups. KO identifiers were mapped to KEGG pathways, and the number of KO genes per pathway was counted for each genome.

The analysis included eight groups: the two *S. chromogenes* strains characterized in this study (4S77 and 4S90); a composite group of six reference *S. chromogenes* genomes (17A, DSM 20454, 1401, IVB6199, IVB6200, and 76ME); and five comparison strains representing ecological and pathogenic diversity: *S. aureus* USA300_FPR3757 (a hypervirulent MRSA strain), *S. aureus* RF122 (a bovine mastitis-associated strain), *S. epidermidis* O47 (a biofilm-forming nosocomial pathogen) [79], *S. epidermidis* ATCC 12228 (a non-pathogenic strain), and *S. carnosus* TM300 (a GRAS-classified starter culture) [18].

KO gene counts per pathway and pathway completeness were analyzed and visualized as clustered heatmaps using Python (v3.13.3) and Seaborn (clustermap, v0.12.2) [80]. For KO abundance, raw counts were used, while completeness was calculated by normalizing each genome’s KO count per pathway to the total number of KO entries defined for that pathway in KEGG (i.e., completeness = observed KOs / total KOs for a pathway). In both cases, hierarchical clustering was performed using Euclidean distance and Ward’s linkage, grouping genomes according to similarity in KO distribution or completeness profiles across pathways. To complement this, pairwise metabolic similarity was also assessed using Jaccard indices calculated from binary presence/absence data. For pathway-level comparisons, a pathway was considered present in a genome if it contained at least one associated KO gene; for KO-level comparisons, presence was determined per individual KO identifier. For each pair of genomes, the Jaccard index was calculated as the size of the intersection divided by the size of the union of the sets of present elements (pathways or KOs), yielding a value between 0 (no shared features) and 1 (identical profiles). These similarity matrices were visualized as clustered heatmaps using hierarchical clustering with average linkage, allowing for topology-based comparisons of metabolic repertoire.

An UpSet plot was also generated to visualize shared and unique pathway distributions among the groups. A binary presence/absence matrix was used as input and processed with the upsetplot library in Python. All analyses and visualizations were conducted in a Jupyter Notebook using the Anaconda 3-2024.10-1 distribution.

### *In silico* genome analyses

Antiviral defense systems were identified using PADLOC-DB v2.0.0 [81], CRISPRCasFinder [82], and CRISPRone [83], while TA loci were predicted using TAfinder 2.0 available through the TADB 3.0 platform [84].

Putative AMR genes were identified using ResFinder v4.7.2 [85] and the Resistance Gene Identifier (RGI, v6.0.3) from the Comprehensive Antibiotic Resistance Database (CARD v4.0.0) [86]. All predictions were manually reviewed to confirm functional relevance, and false positives, such as alanine racemase misidentified as *vanT* by CARD, were excluded from the final results. To assess potential resistance mechanisms against AMPs, all genomes were screened using BLASTx with reference amino acid sequences of known AMP defense proteins. This included cell envelope-modifying systems (*mprF*, *dltABCD* operon), regulatory and transporter modules (*braSR/braDE*, *graRS/vraFG*, *vraDEH*), the *icaADBC* operon linked to biofilm-associated protection, the *capBCAD* operon involved in PGA capsule synthesis, and proteases known to degrade AMPs (*aur*, *sspA*, *sak*).

Putative VFs were first screened using the VFanalyzer tool from the Virulence Factor Database (VFDB) with default parameters [87]. To complement this, a parallel BLASTx search was carried out using a custom set of staphylococcal VF protein sequences compiled primarily from Naushad et al. [51], along with several additional factors identified through literature review [55, 88–91]. Hits were filtered with a minimum threshold of 30% identity and 30% alignment coverage, and all candidates were manually validated using BLASTp to confirm functional relevance. For instance, a gene initially flagged as *sbnA* was excluded after protein-level analysis showed it encoded cysteine synthase, a distant SbnA homolog. The complete list of reference VF sequences used for screening is provided in supplementary S1 File.

To characterize the regulatory landscape of *S. chromogenes*, TCSs were identified using Predicted Prokaryotic Regulatory Proteins (P2RP) [92] and validated through BLASTx against a curated set of experimentally characterized *S. aureus* TCSs described by Bleul et al. [27].

ISs were identified using ISfinder with BLASTN v2.2.31+ (default parameters) [93] and MobileElementFinder (v1.0.3, MGEdb v1.0.2) with a minimum of 30% identity and coverage, and a maximum truncation of 30% [94]. Putative composite Tns and Tn3-family transposable elements were identified using Composite Transposon Finder and Tn3 Transposon Finder, respectively, under default parameters [95]. Genomic islands were predicted using

IslandViewer 4 [96], prophage regions were identified with PHASTEST [97], and SCCmecFinder v1.2 [98] was used to screen for SCCmec elements. Plasmid-encoded operons were predicted using Operon-mapper with default parameters [99]. Plasmid replicon types were identified using PlasmidFinder v2.1 (database v2023-01-18) with the Gram-positive database, a minimum identity threshold of 50%, and a minimum coverage of 20% [100].

BGCs were identified using the bacterial version of antiSMASH v7.0 [101], BAGEL4 [102], and BACTIBASE [103].

Phobius was used to predict signal peptides and TM topology of proteins, based on combined hidden Markov models for both features [104].

All amino acid sequence alignments were performed using the Constraint-Based Multiple Alignment Tool (COBALT) from NCBI, which aligns sequences based on conserved domain information and sequence similarity.

Comparative gene cluster diagrams were generated using clinker through the CompArative GEne Cluster Analysis Toolbox (CAGECAT) platform.

### Antibiotic susceptibility assays

MIC values for norfloxacin and lincomycin were determined using the broth microdilution method in cation-adjusted Mueller-Hinton broth (Merck Co., Darmstadt, Germany), with antibiotic concentrations ranging from 32 to 0.0625 µg/mL. Bacterial inocula were added to achieve a final concentration of 5×10⁵–1×10⁶ CFU/mL per well, and plates were incubated at 37 °C with shaking for 18 h. MICs were defined as the lowest concentration with no visible growth and were assessed in triplicate. Clindamycin susceptibility was evaluated separately using an MIC Test Strip (Liofilchem, Roseto degli Abruzzi, Italy) on Mueller-Hinton agar (Merck Co.) inoculated to an OD_600_ of 0.1.

### Phenotypic detection of extracellular enzyme activity and biofilm formation

To assess extracellular enzyme activities, overnight cultures were adjusted to an OD_600_ of 0.1, and 10 µL was spotted onto assay plates. Lipase activity was tested on tryptic soy agar (TSA, Merck Co.) supplemented with 1% Tween 20, hemolysis on TSA sheep blood agar (Thermo Fisher Scientific, Germany), and nuclease activity on DNase test agar (Difco, Ausburg, Germany). Plates for lipase and hemolysis were incubated overnight at 37 °C, followed by 24 h at 4 °C before evaluating halo formation. For DNase activity, plates were incubated at 37 °C overnight and then flooded with 1 N hydrochloric acid to visualize zones of clearance. Biofilm formation was assessed by streaking fresh colonies onto Congo red agar and incubating at 37 °C for 24 h. Biofilm-positive colonies were identified based on black or dark-red pigmentation on the agar surface [105].

### MP1 isolation and characterization

Strains 4S77 and 4S90 were cultivated in 1 L of tryptic soy broth at 37 °C for 24 hours. Following centrifugation, the culture SN was extracted with ethyl acetate (EtOAc) at a ratio of 5:1 (SN:EtOAc) at room temperature with continuous agitation. The aqueous phase exhibited no detectable antimicrobial activity and was consequently excluded from further purification. The EtOAc phase was evaporated at 50 °C using a rotary evaporator (R-80, Büchi, Flawil, Switzerland) connected to a vacuum pump (V-180, Büchi), and the resulting residue was dissolved in 5 mL of pre-warmed dimethyl sulfoxide. This crude extract was applied to a CHROMABOND^®^ RS40 C18 cartridge (43 g; Macherey-Nagel, Düren, Germany) operated via an ÄKTA Pure 25 chromatography system (GE Healthcare, Uppsala, Sweden). Elution was carried out using a stepwise gradient of 0–60% buffer B (acetonitrile containing 0.1% trifluoroacetic acid) against buffer A (0.1% trifluoroacetic acid in water) at a flow rate of 10 mL/min. MP1 eluted at 60% buffer B, and the relevant fractions were pooled and assessed for purity by RP-HPLC using a Poroshell 120 EC-C18 column (Agilent Technologies; 4.6×150 mm, 2.7 µm). The purified compound was lyophilized for 24 hours and subsequently reanalyzed by RP-HPLC to confirm its purity.

To confirm the identity of the antimicrobial compound as MP1, LC-MS was performed using an Agilent 1200 HPLC-MS system equipped with a 1200 diode array detector (10 mm standard flow cell) and an LC/MSD Ultra Trap System XCT 6330 (Agilent, Waldbronn, Germany). A sample volume of 2.5 μL was injected onto a Reprosil C18 column (100 Å, 2 mm ID) with a 10 Å, 2 mm ID precolumn (Dr. Maisch GmbH, Ammerbuch, Germany), operated at a flow rate of 400 μL/min and a column temperature of 40 °C. Analytes were separated using a 20-minute gradient from 100% solvent A (water with 0.1% formic acid) to 100% solvent B (acetonitrile with 0.06% formic acid), followed by a 5-minute elution at 100% B. Mass spectrometry was carried out in alternating positive and negative electrospray ionisation modes, with a capillary voltage of 3.5 kV and an ultra-scan range up to *m/z* 2000. Data were analyzed using 6300 Series Trap Control software (version 6.1, Agilent).

The antimicrobial activity of the crude extract, chromatographic fractions, and purified MP1 was evaluated using a well diffusion assay against *S. aureus* USA300 JE2. In brief, 50 µL of each sample was loaded into wells on soft TSA (0.6% agar) plates seeded with 1×10⁶ CFU/mL of the test strain, and zones of inhibition were recorded after overnight incubation at 37 °C.

### Synthetic peptides and antimicrobial activity assay

Two peptides encoded by the unidentified RiPP-like gene cluster in *S. chromogenes*, comprising 42 and 51 amino acids, were synthesized with and without their predicted N-terminal leader sequences. Peptides were synthesized at >95% purity by APeptide Co., Ltd (Shanghai, China). Antimicrobial activity was evaluated using a well diffusion assay against *Micrococcus luteus* ATCC 4698. Synthetic peptides were tested individually and in equimolar combinations at a final concentration of 100 µM. Peptide A from aureocin A70 served as a positive control.

### Distribution Analysis of the MP1 Gene Cluster

To assess the distribution of the MP1 BGC across the genus *Staphylococcus*, a dataset of 2,939 unique complete genomes was compiled from NCBI (retrieved on 11 December 2024), excluding duplicate assemblies of the same strain (**S14 Table**). Homologs of the MP1 precursor peptide from *S. chromogenes* strains 4S77 and 4S90 were identified using a tBLASTn search in CLC Genomics Workbench 20.0 (Qiagen, Hilden, Germany), with the following parameters: E-value threshold of 0.01, BLOSUM62 matrix, no masking of lower-case sequences, and genetic code 11 (bacterial). Search results were filtered manually to identify hits with conserved synteny and complete cluster content.

To further investigate whether the MP1 BGC had been previously reported in *S. chromogenes*, an independent BLASTp search was conducted on 5 December 2025 against the NCBI non-redundant protein database (nr). The MP1 precursor peptide from *S. chromogenes* strains 4S77 and 4S90 was used as the query, with the search restricted to the taxon *S. chromogenes* (taxid:46126).

## Data availability

The genome sequences of *S. chromogenes* strains 4S77 and 4S90 were deposited in GenBank under the accession numbers CP195074–CP195075 (4S77) and CP195066– CP195068 (4S90), respectively, under BioProject PRJNA1273467.

## Acknowledgment

We would like to kindly thank Dr. Libera Lo Presti for her valuable comments and suggestions on scientific writing.

## Funding

This work was supported by the Deutsche Forschungsgemeinschaft (DFG, German Research Foundation) Germany’s Excellence Strategy—EXC 2124—390838134 ‘Controlling Microbes to Fight Infections’ (CMFI). We acknowledge the support by Open Access Publishing Fund of University of Tübingen.

## Abbreviations

AAA: aromatic amino acid
AMP: antimicrobial peptide
AMR: antimicrobial resistance
BCAA: branched-chain amino acid
BCFA: branched-chain fatty acids
BGC: biosynthetic gene cluster
CsoR: copper-sensing transcriptional repressor
CsoZ: putative copper chaperone
EtOAc: ethyl acetate
GEH: glycerol ester hydrolase
GG: glycine-glycine
GI: genomic island
GRAS: generally recognized as safe
HEPN: higher eukaryotes and prokaryotes nucleotide-binding
IS: insertion sequence
KO: KEGG ortholog
MP1: micrococcin P1
MRSA: methicillin-resistant Staphylococcus aureus
msr: macrolide-streptogramin resistance
NAS: non-aureus staphylococci
PDC: phage defense candidate
PGA: poly-?-glutamic acid
RiPP: ribosomally synthesized and post-translationally modified peptide
RM: restriction-modification
RP-HPLC: reverse-phased high-performance liquid chromatography
S.: Staphylococcus
SCC: staphylococcal cassette chromosome
SN: supernatant
TA: toxin-antitoxin
TCS: two-component system
TM: transmembrane
Tn: transposon
VF: virulence factor.

